# Gene regulatory networks linked to GABA signalling emerge as relevant for glioblastoma pathogenesis

**DOI:** 10.1101/2025.04.30.651564

**Authors:** Itzel Nissen, Soran Dakhel, Chaitali Chakraborty, Amalie Holm Nygaard, Agnete Kirkeby, Andreas Hörnblad, Cemal Erdem, Silvia Remeseiro

## Abstract

Glioblastoma (GB) is the most common and aggressive adult brain tumor. Recent evidence shows that GB cells form glutamatergic synapses with neurons, promoting tumor growth and invasion, yet the gene regulatory networks (GRNs) underlying this interaction remain underexplored. GRNs coordinate gene expression through interactions among transcription factors (TFs), enhancers, and promoters, and are often disrupted in cancer due to epigenetic and chromatin architecture changes. To identify GRNs involved in GB pathogenesis, we applied the machine learning-based MOBILE (Multi-Omics Binary Integration via Lasso Ensembles) pipeline to integrate multi-omics data. GABA-signalling emerged as a previously unrecognized contributor to GB, involving relevant TFs as ARX, GSX2 and DLX family, key regulators of GABAergic interneuron development. Co-culture assays further demonstrated that GABAergic input promotes GB proliferation through non-synaptic mechanisms likely involving metabolic or paracrine interactions. Our findings reveal novel GRNs in GB, positioning GABA signalling as a potential therapeutic target to disrupt neuron-glioma interactions.

**Significance:** This study uncovers novel gene regulatory networks driving glioblastoma progression through integration of multi-omics data with machine-learning tools. By identifying GABAergic signalling as a previously unrecognized contributor to tumour growth, it reveals new transcriptional regulators, providing valuable insights into neuron-glioma interactions. These findings open new therapeutic avenues for brain-cancer treatment.

**GRAPHICAL ABSTRAC:** 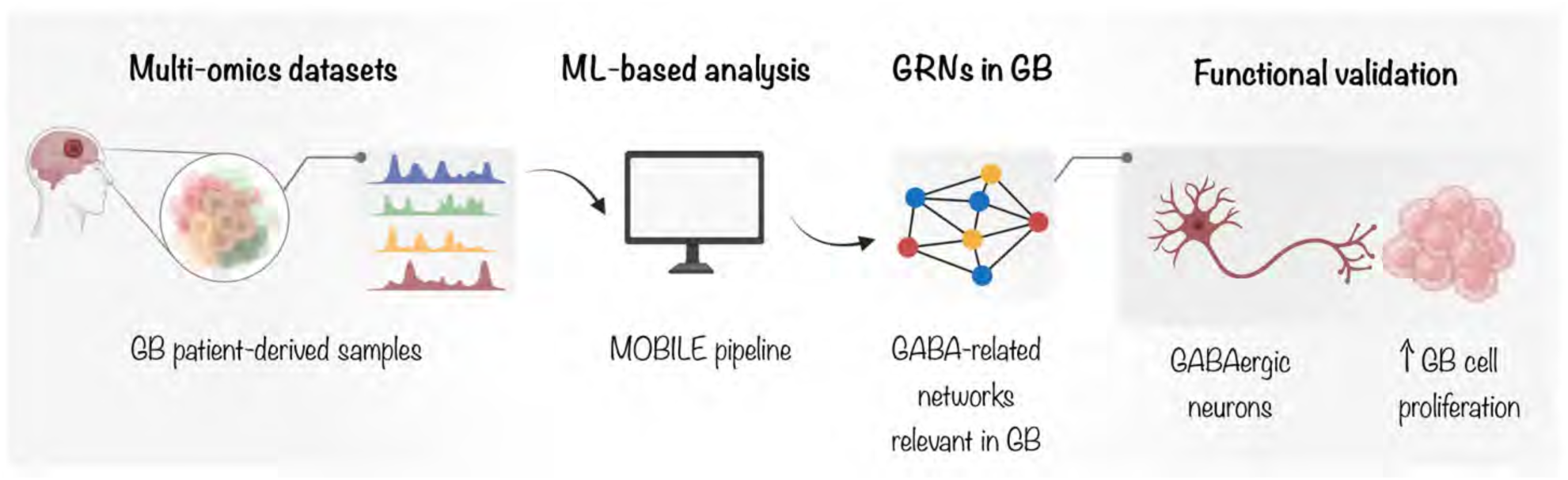

Integration of multi-omics data from glioblastoma (GB) patients identified networks related to GABA signalling as relevant for GB pathogenesis. In co-culture assays, GABAergic interneurons increase the proliferation of GB cells.

## INTRODUCTION

Glioblastoma (GB) is the most common and aggressive primary brain tumour in adults. Its poor prognosis and low survival rate (5-year survival ∼4%), rapid growth and high invasive capacity, alongside its resistance to standard therapies make GB a formidable clinical challenge [1,2]. GB tumours exhibit high molecular heterogeneity, reflected by the presence of multiple cellular subtypes with distinct transcriptional and epigenetic landscapes [3–6]. Recent discoveries have shown that GB cells can establish *bona fide* synapses with neurons present in the tumour surroundings. These are glutamatergic synapses that relay electrical signals to glioma cells via AMPA receptors [7,8]. Neuron-to-glioma synapses are believed to foster a favourable microenvironment, driving tumour progression by promoting glioma cell growth and hijacking neural mechanisms of invasion [7–10]. However, a key yet underexplored question is how glioma cells decode these neuronal signals at the genomic level. In this context, our group recently identified gene regulatory networks (GRNs) through which this neuron-to-glioma synaptic communication operates, establishing a mechanistic link between neural input and tumour growth [11].

GB genomes and epigenomes are extensively rewired, leading to altered GRNs that drive transcriptional programs sustaining its aggressiveness and malignancy [11–17]. GRNs are critical for maintaining cellular identity and function by precisely coordinating gene expression through interactions among transcription factors (TFs), enhancers, and gene promoters. Enhancers are *cis*-regulatory elements critical for initiating and modulating gene transcription in a temporal- and tissue-specific manner. Enhancer activity is tightly connected to epigenetic states, which are defined by several features including the frequency of observation of specific histone modifications and other chromatin features [18]. Thus, enhancers exhibit different states (i.e., active, primed, poised or silenced) emerging from these combinations of histone modifications, which subsequently determine how regulatory elements impact transcription at a given time [19–20]. While H3K27me3 is a repressive histone modification, H3K27ac marks active enhancers; and although H3K4me3 is more prominent at active promoters, it is also present at active enhancers [19–22]. Importantly, enhancers can be located very far away from their target genes in the linear genome and can therefore act across large genomic distances [23]. This is facilitated by the 3D organization of the genome, which brings enhancers and promoters in physical proximity, allowing them to establish regulatory interactions [24–26]. Moreover, enhancers serve as platforms to recruit transcription factors (TFs) through short, specific DNA sequence motifs to regulate transcription [27]. Sustaining the intricate gene expression programs necessary to determine cellular identity and function therefore requires complex GRNs, including enhancer-promoter regulatory interactions and the function of specific TFs. Additionally, genome-wide enhancer mapping has revealed that many disease-associated genetic variants are located within non-coding regulatory regions, particularly enhancers, highlighting their relevance in the pathogenesis of various diseases, including cancer [28,29]. Cancer cells frequently exhibit extensive reorganization of chromatin architecture and enhancer-promoter interactions, leading to dysregulated transcriptional networks that support tumourigenesis [30]. In this context, disruptions in GRNs often result from mutations, epigenetic alterations, and changes in enhancer-promoter connectivity. Disruption of insulated neighbourhoods, enhancer hijacking or somatic mutations in regulatory elements, are therefore different mechanisms that can lead to the acquisition of oncogenic enhancers that in turn activate proto-oncogenes, leading to tumourigenesis [31–33]. Understanding these shifts in regulatory landscapes is essential for understanding many molecular mechanisms underlying cancer.

Using a comprehensive multi-omics approach, we previously uncovered a rewiring of the enhancer-promoter interactome and regulatory landscape that sustains transcriptional programs allowing glioma cells to integrate neural input into a proliferative response [11]. SMAD3 and PITX1 were identified as the major TFs in a GRN supporting specific transcriptional programs related to glutamatergic signalling, axon guidance and axonogenesis [11]. Moreover, in functional assays, we demonstrated that targeting specific TFs and their associated networks can modulate tumour growth [11], supporting that epigenetic regulators of this neuron-to-glioma axis may represent clinically relevant therapeutic targets. Nonetheless, accurately capturing the functional and biologically relevant epigenomic and transcriptomic changes in cancer remains a challenge. This has driven the development of advanced computational approaches that integrate complex, multi-layered epigenetic and transcriptional data to reveal the key networks that are functionally relevant in cancer [34–36].

In this study, we applied a machine learning (ML)-based method [37] to a multi-omics dataset-including transcriptome profiles, chromatin accessibility and histone modifications-that we obtained from a panel of 15 patient-derived GB cell lines, representing the four GB expression subtypes, alongside normal human astrocytes. Our analyses uncovered novel GRNs and pointed to additional TFs relevant for GB pathogenesis. GABA-related signalling emerged as a previously unrecognized driver of GB pathogenesis, and TFs such as ARX, GSX2 and DLX family, among others, were identified as relevant regulators in the context of neuron-glioma communication. Moreover, we functionally demonstrate that, in co-culture assays, not only glutamatergic neurons but also GABAergic interneurons can promote the proliferation of GB cells. Our findings highlight the relevance of new GRNs associated to GABA signalling in neuron-glioma interactions, presenting promising new therapeutic targets for highly treatment-resistant cancers such as glioblastoma.

## RESULTS

### ML-based approaches identify new gene regulatory networks relevant in glioblastoma

Advanced computational approaches provide powerful tools for analysing multi-omics data, uncovering intricate patterns and relationships that often go undetected by traditional bioinformatics methods. In a recent study, our group generated a multi-omics dataset from 15 patient-derived glioblastoma (GB) cell lines, alongside normal astrocytes as control, and identified the major gene regulatory networks (GRNs) underlying neuron-to-glioma synaptic communication in GB [11] . Here, we applied a machine learning (ML)-based approach to this multi-omics dataset (Fig. 1A), revealing additional networks and mechanisms governing this complex disease. For this purpose, we used MOBILE (Multi-Omics Binary Integration via Lasso Ensembles), a pipeline designed to integrate multi-omics datasets, allowing the inference of context-specific interactions and regulatory pathways [37]. The omics data here includes transcriptome profiles (RNA-seq), chromatin accessibility (ATAC-seq) and ChIP-seq for histone modifications H3K27ac (active enhancers), H3K4me3 (active promoters) and H3K27me3 (repressive regions). Using the MOBILE pipeline, we applied the Lasso module to integrate the omics datasets pairwise. We paired ATAC-seq with RNA-seq (ATAC→RNA), and ChIP-seq for H3K27ac (H3K27ac→RNA), H3K27me3 (H3K27me3→RNA), and H3K4me3 (H3K4me3→RNA) with RNA-seq, such that X→RNA refers to the effect of peak X on the expression of a certain gene identified by RNA-seq, where X represents either an ATAC peak or ChIP peak for any of the histone modifications. This analysis includes both the 15 GB lines and normal astrocytes, and it results in a Lasso coefficient matrix, which contains association coefficients between analytes from the two input matrices. The mean values of these coefficients populate the FULL

**Figure 1.**
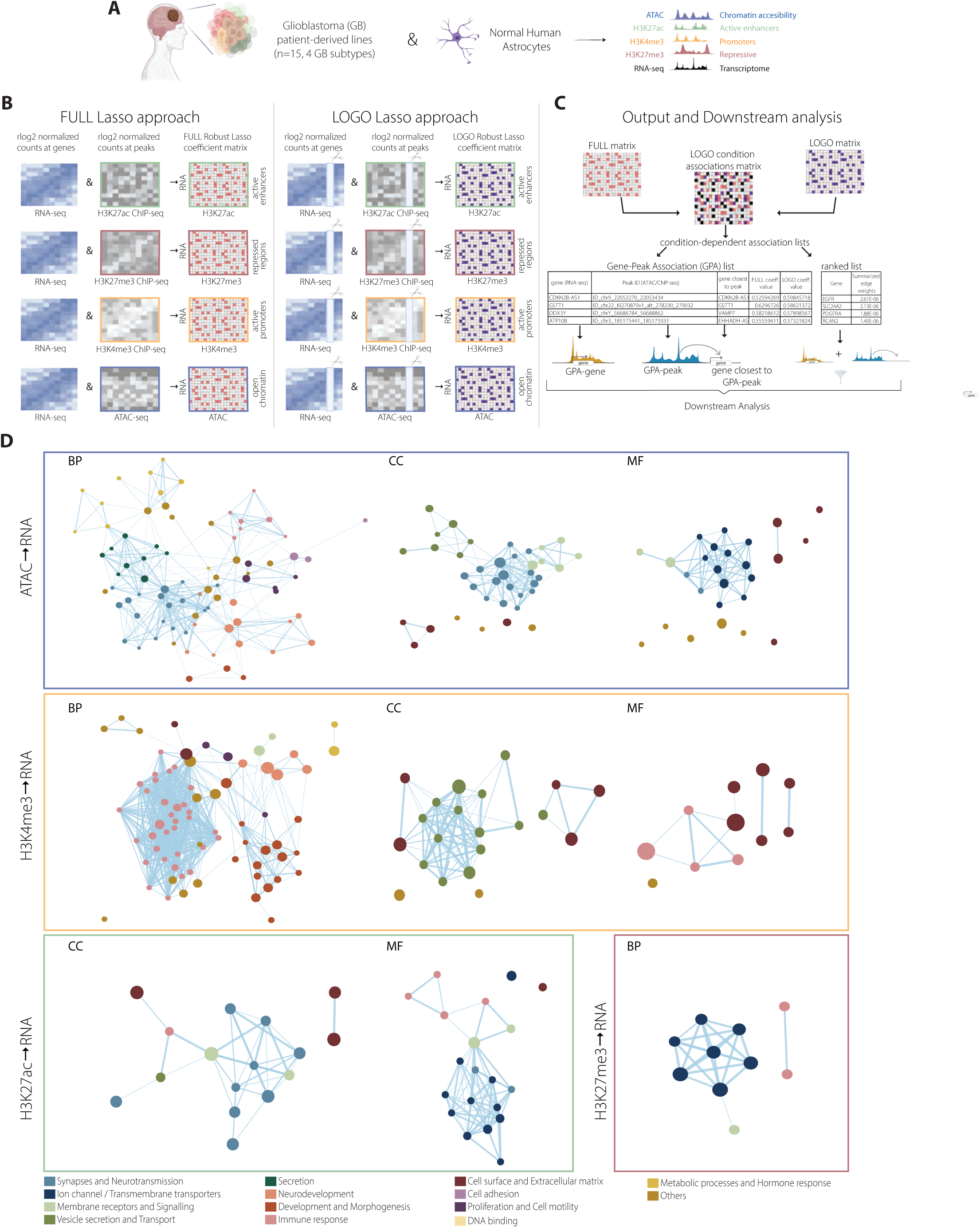
ML-based approach identifies context-specific networks relevant in GB. **(A-C)** Workflow of the MOBILE pipeline. Multi-omics datasets from 15 patient-derived GB lines and normal human astrocytes from Chakraborty et al., 2023 [7] were used (A). The Lasso module integrates omics datasets in pairs, producing a coefficient matrix containing the association coefficients between analytes of the two input matrices. Mean values of these coefficients form the FULL Robust Lasso Coefficient Matrix. Excluding the normal human astrocytes, the LOGO (leave-one-group-out) module is then run to generate LOGO Robust Lasso Coefficient Matrices (B). Comparing the FULL and LOGO matrices revealed GB-specific associations, yielding a condition-dependent associations gene list and a ranked list that were then used for further downstream analysis (C). **(D)** Identification of GB-associated networks showing enrichment for synapses and neurotransmission, immune response, proliferation and cell motility, among others.

Robust Lasso Coefficient Matrix (Fig. 1B, left). Subsequently, human astrocytes were excluded from the training input, and the LOGO (leave-one-group-out) module was implemented, generating the LOGO Robust Lasso Coefficient Matrices (Fig. 1B, right). By comparing the FULL association matrix to the LOGO matrix, we identified associations specifically dependent on the exclusion of astrocytes, and therefore GB-specific (Fig. 1C). This analysis yielded: i) condition-dependent associations lists, and ii) ranked lists, where the genes in the associations list (with lasso coefficient with a magnitude larger than 0.01) are ordered based on the strength of the relationships (edges) between entities (nodes) in the inferred network (Supplementary Data 1). Each associations list compiles gene-peak associations (GPAs), reflecting that a certain ATAC/ChIP peak is associated to the expression of a certain gene in GB. This list therefore contains two distinct gene sets: i) genes in gene-peak associations, hereby named GPA-genes, and ii) genes closest to GPA-peaks. Both the associations lists and the ranked-genes lists were further examined in downstream analyses.

### Novel GRNs in GB are related to GABA-signalling, ECM and immune response

Given the power of ML-based approaches to uncover intricate patterns and interactions that often escape other methods, we anticipated that MOBILE would identify novel GRNs linked to pathways not previously recognized in GB pathogenesis. To determine which biological processes were captured by this analysis, we conducted gene set enrichment analysis (GSEA) on the GB-specific ranked lists (i.e., LOGO-condition). Significantly enriched terms and pathways (p<0.05) were identified for biological processes (BP), molecular function (MF), and cellular component (CC) for each pairwise comparison, i.e., ATAC→RNA, H3K27ac→RNA, H3K27me3→RNA, and H3K4me3→RNA (Fig. 1D and Supplementary Figs. 1-3). Some of the identified pathways were common to all four analyses. Among the most represented terms were those related to neural functions: synapses, neurotransmission, ion channels and transmembrane transporters, vesicle secretion and transport, as well as neurodevelopment. This aligns with previous findings highlighting the significance of neuron-glioma interactions in GB [7,8,38,39] and the first identification of GRNs underlying neuron-to-glioma synapses in GB [11]. Additional enriched terms were linked to immune response, extracellular matrix (ECM) composition and remodelling, as well as cell motility and proliferation. Similarly, prior studies have shown that glioblastomas actively remodel the ECM to promote invasion and evade immune detection [40–44].

To explore these pathways in deeper detail, we performed Gene Ontology (GO) analysis on the ranked-list genes from each GB-specific pairwise comparison (Supplementary Figs. 4 and 5). The top significantly enriched GO terms were associated to synapses, including glutamatergic and GABAergic synapses, ion channel activity, ECM structure and organization, MHC complexes, and immune response. This agrees with findings on the mechanisms used by GB to remodel the ECM in its own benefit in order to promote recurrence and tumour progression, and to escape the immune system [40,43]. MHC I levels are commonly reduced in glioblastoma, favouring evasion of CD8+ T cell recognition, and weakening NK cell responses by reducing their ability to identify and target GB cells [45]. Networks related to MHC protein complexes, and complex binding, were significantly enriched in our data (Supplementary Figs. 4 and 5). Altogether, this suggests that the chromatin accessibility and epigenetic changes occurring in GB could influence immune recognition, tumour immunogenicity and ECM remodelling through networks hereby identified. Importantly, we also identified further pathways related to neurotransmission, in particular networks linked to both glutamatergic and GABAergic signalling (Supplementary Figs. 4 and 5). Glutamatergic synapses between neurons and GB cells were described as drivers of tumour progression [7,8], and our group recently identified the major TFs and GRNs underlying this neuron-to-glioma synaptic communication [11]. Diffuse hemispheric gliomas (DHG-H3G34 mutant) display a GABAergic interneuronal lineage identity [46], and GABAergic neuron-to-glioma synapses were recently observed in diffuse midline gliomas (DMG), a type of paediatric high-grade glioma [47]. However, to date, there is no experimental confirmation of GABAergic synaptic connections between neurons and GB cells. Our computational analysis consistently captured GABAergic signalling as a significantly enriched pathway, suggesting that GABAergic signalling might play an important role in GB pathogenesis.

Additionally, comparing the ranked-list genes generated by MOBILE with the set of 497 differentially expressed genes (DEGs) that we previously identified with DESeq2 analysis [11], we observed a substantial overlap, i.e., 20% of the 497 DEGs overlap with genes in the ranked list (Supplementary Fig. 6). Both approaches highlight neural function-related processes as the most prominent and relevant to GB pathogenesis (Supplementary Fig. 6). However, compared to the number of DEGs detected through standard differential expression analysis [11], MOBILE captured a substantially larger set of genes as relevant for GB pathogenesis. Moreover, the genes identified by MOBILE are associated to changes in chromatin accessibility and histone modifications, providing a more comprehensive inference through data integration.

We then examined our data more closely, accounting for differences in active (open chromatin and/or active histone marks) *vs* inactive (presence of H3K27me3) epigenetic states. By overlapping the genes from the four ranked lists obtained through pairwise comparisons, we retrieved sets of genes linked to either active or inactive regions (Fig. 2A). GO analysis of genes associated to inactive regions revealed a significant enrichment for terms related to negative regulation of cell migration, cell cycle and angiogenesis (Fig. 2B). The H3K27me3-dependent inactivation of negative regulators of cell migration and cell cycle is consistent with the high levels of invasion and proliferation characteristic of GB. In contrast, genes associated with active regions were enriched for processes such as neurotransmission, synapse organization - including GABAergic and glutamatergic synapses-as well as axonogenesis and axon guidance (Fig. 2C), suggesting that these processes are stimulated in GB.

**Figure 2.**
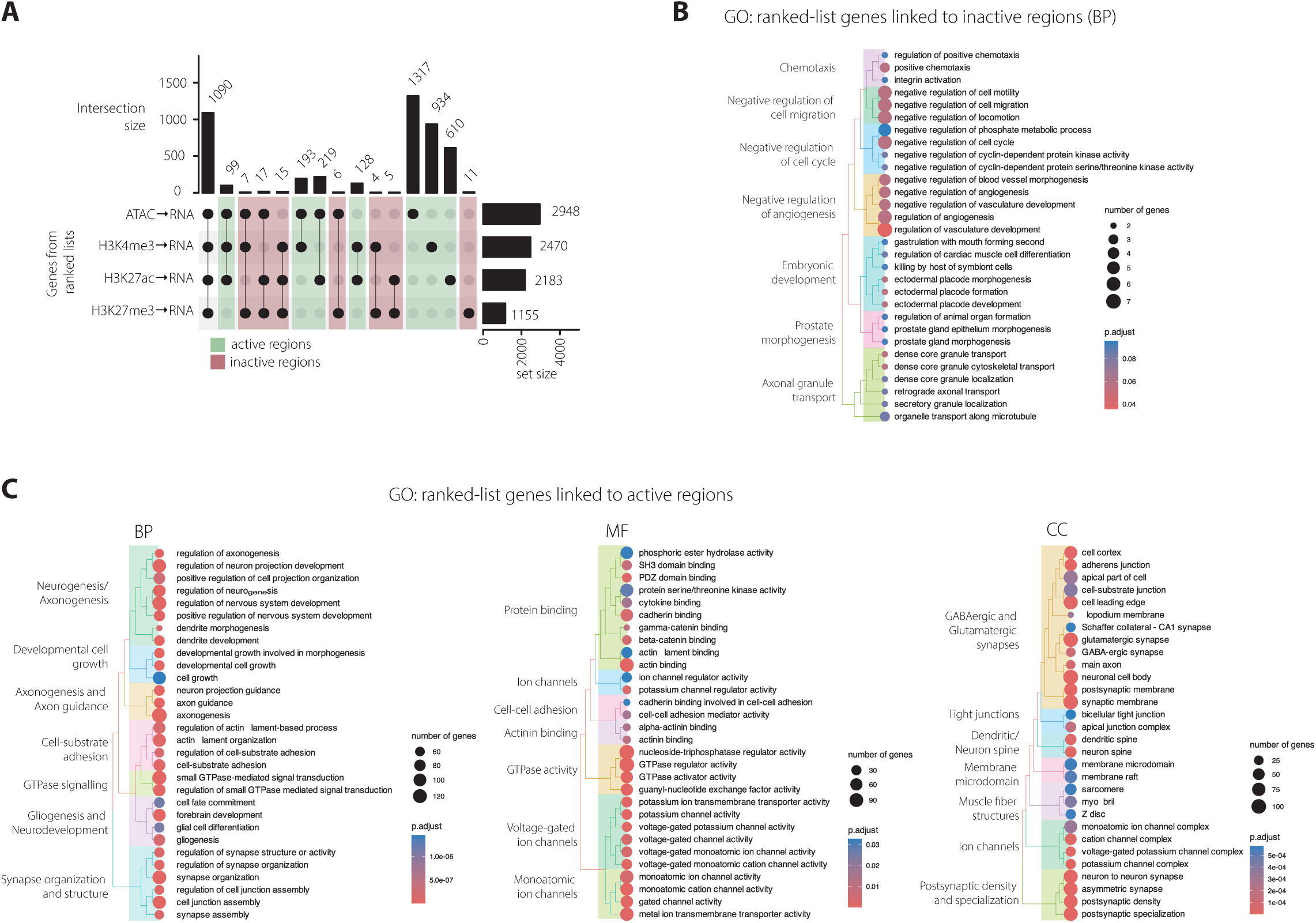
Association of GB-specific networks and active/inactive states. **(A)** Intersection of the four ranked lists segregates genes as being associated to either active regions (only open chromatin and/or H3K27ac, H3K4me3) or inactive/repressed regions (presence of H3K27me3). **(B-C)** GO analysis of ranked-list genes linked to inactive regions shows enrichment for GO terms related to negative regulation of: cell migration, cell cycle and angiogenesis (B), while genes linked to active regions are associated to synapses, including GABAergic and glutamatergic synapses, neurotransmission, axon guidance and cell adhesion (C).

While capturing previously known pathways, the integration of multi-omics data using MOBILE also identified novel GRNs relevant for GB pathogenesis, including the previously unrecognized role of GABAergic signalling as a driver of GB.

### MOBILE detects relevant expression changes not identified by standard differential expression analysis

Differential expression analysis identifies genes with significant changes in expression levels between different conditions, and it is typically based on statistical comparison of normalized gene expression data obtained often from RNA-sequencing. Instead, the MOBILE pipeline captures intricate patterns and relationships by integrating multi-omics datasets. In this study, MOBILE has identified genes and their associations with changes in chromatin accessibility and histone modifications, highlighting their relevance within specific networks.

Our analysis points to GO terms related to immune function, extracellular matrix (ECM), as well as GABA and glutamate signalling as the most prominent for GB pathogenesis (Fig. 1D, 2B-C and Supplementary Fig. 1-5). Next, we retrieved the particular genes within these GO terms and more closely examined their gene expression patterns across the 15 GB lines relative to normal human astrocytes. MOBILE identified 30 genes related to MHC-complex function as significantly relevant for GB-specific networks, despite subtle changes in expression when comparing the GB lines to astrocytes (Fig. 3A, C). Out of those, only *HLA-A* and *HLA-B* genes had been identified in our previous study as significantly downregulated in all 15 GB lines *vs* normal astrocytes (497 DEGs in [11]). Tumour cells evade immune detection by downregulating HLA-A and HLA-B expression [48] and, in colorectal cancer, high HLA-B/C expression is an independent predictor of favourable prognosis [49]. In addition to *HLA-A* and *HLA-B,* MOBILE captured additional *HLA* genes (e.g., *HLA-DRA, -DRB1/5, -F, -E, -DOA*), and other immune-related genes like *TAP1, TAP2* and *CD4.* On other hand, 237 genes related to ECM functions were identified as significantly relevant for GB networks (Fig. 3B, C), out of which only 11 were identified by DESeq2 in our previous study [11]. Among those, we found ECM modulators *BCAN*, *LOXL4, L1CAM*, and *FOXC2* which support ECM remodelling and cancer progression. BCAN activates EGFR/MAPK signalling and fibronectin production in glioma cells, enhancing their migratory and invasive properties, making it a target under study for GB treatment [50]. While LOXL4 promotes ECM remodelling and tumour cell migration through tissue stiffening, a common feature in invasive cancers [51], L1CAM (L1 cell adhesion molecule) promotes cell migration [52] and FOXC2 is critical for epithelial-mesenchymal transition [53]. Additional genes identified by MOBILE, which went undetected previously, include seven genes encoding matrix metalloproteinases (*MMP*s), 28 genes from collagen family (*COL*), *FN1*, *TGFBI* and *SPARCL1*, all of which have been prominently implicated in cancer biology [54–58].

**Figure 3.**
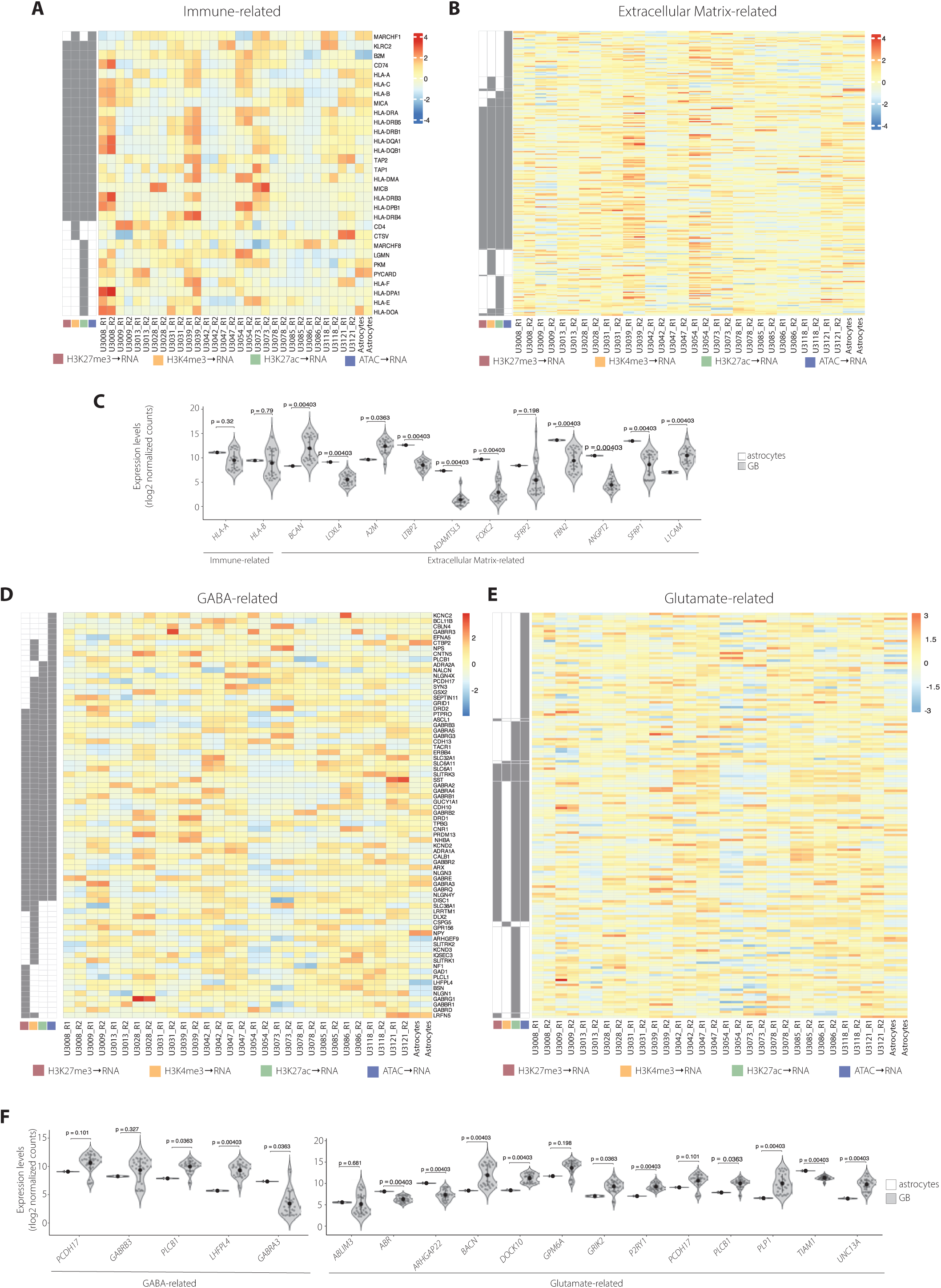
Changes in the expression of genes within immune-, ECM-, GABA- and glutamate-related networks identified as relevant in GB. **(A-B)** Immune-related (A) and ECM-related (B) pathways were identified by MOBILE as significant in GB pathogenesis, despite the fact that genes within these networks display moderate gene expression changes, not detectable by standard gene expression analysis. **(C)** Expression of genes in the networks identified in A-B that overlap with the 497 DEGs in 15 GB lines vs astrocytes from Chakraborty et al., 2023 [7]. **(D-E)** GABA-related (D) and glutamate-related (E) pathways were identified by MOBILE as significant in GB pathogenesis, despite genes within these networks display moderate gene expression changes, not detectable by standard gene expression analysis. **(F)** Expression of genes in the networks identified in D-E that overlap with the 497 DEGs in 15 GB lines vs astrocytes from Chakraborty et al., 2023 [7].

Similarly, we examined in closer detail the genes contributing to GABAergic and glutamatergic signalling, pathways identified as relevant for GB pathogenesis (Fig. 3D-F). Despite subtle changes in expression in GB cells relative to astrocytes, MOBILE could identify 74 genes related to GABA-signalling (Fig. 3D) and 165 related to glutamate-signalling (Fig. 3E) as relevant in GB malignant networks. Out of those, only 5 and 13, respectively (Fig. 3F), had been identified by DESeq2 in our previous work [11]. Notably, among the genes detected by MOBILE are those encoding GABA receptors (e.g., *GABRA2/3/4, GABRB1/2/3, GABRE, GABRG1, etc.*), ion channels involved in neurotransmission (e.g., *KCNC2, KCND2, KCND3*), transcription factors involved in neural differentiation and regulation of GABAergic neurons (e.g., *ARX*, *DLX2*), alongside synaptic adhesion molecules (e.g., neuroligins *NLGN4X, NLGN1*and *NLGN3*) and *GAD1,* encoding the enzyme catalysing the synthesis of GABA. Interestingly, NLGN3 localizes at both excitatory and inhibitory synapses [59,60], and neural activity-regulated secretion of NLGN3 promotes the proliferation of high-grade glioma cells [38,39]. On the other hand, MOBILE also captured genes related to glutamatergic signalling, including those encoding glutamate kainate receptors (e.g., *GRIK2, GRIK4*), AMPA (e.g., *GRIA1, GRIA4*), NMDA (e.g., *GRIN2A*), delta (e.g., *GRID2*) and mGluR (e.g., *GRM7* and *GRM8*) receptors. Noteworthy, genes such as *PLCB1* and *PDCH17* have been shown to modulate both GABAergic and glutamatergic signalling [61,62]. These findings further support the relevance of not only glutamatergic but also GABAergic signalling for GB malignant networks.

While glutamatergic neuron-to-glioma synapses drive tumour progression and invasion in GB and other gliomas [7,8], GABAergic neuron-to-glioma synapses have only been described in pediatric high-grade gliomas [47], but have not been observed in GB to date. In brain metastases, tumour cells adapt a GABAergic phenotype, exploiting inhibitory signals and glutamine metabolism to create a tumour-permissive microenvironment; and importantly, disruption of GABA metabolism with the antiepileptic drug vigabatrin reduces brain metastasis [63]. Our present analysis, which identifies GABA-related networks as key drivers of GB malignancy, together with studies in gliomas and brain metastasis, raises the question of whether GABAergic neuron-to-glioma synapses, or other forms of GABAergic neuron-tumour interactions, are as relevant in GB as they appear to be in high-grade gliomas such as DMGs [47].

### ARX, GSX2 and DLXs identified as TFs relevant for GABAergic-related networks in GB

Having identified networks relevant for GB malignancy, we next examined through which regulators these GRNs might operate. Based on Panther annotations, we found 92 genes encoding TFs among the GPA-genes, while an additional 348 TF-encoding genes were annotated as genes closest to GPA-peaks (Fig. 4A, left and middle, respectively). Among the genes in the ranked lists, 301 genes encoding TFs were identified (Fig. 4A, right). Notably, there are additional relevant genes in the networks coding for histone modifiers and other chromatin binding proteins (Fig. 4A). This suggests that GRNs associated with GB malignancy likely consist of numerous transcription factors and are regulated by a complex interplay between TFs and chromatin-associated proteins.

**Figure 4.**
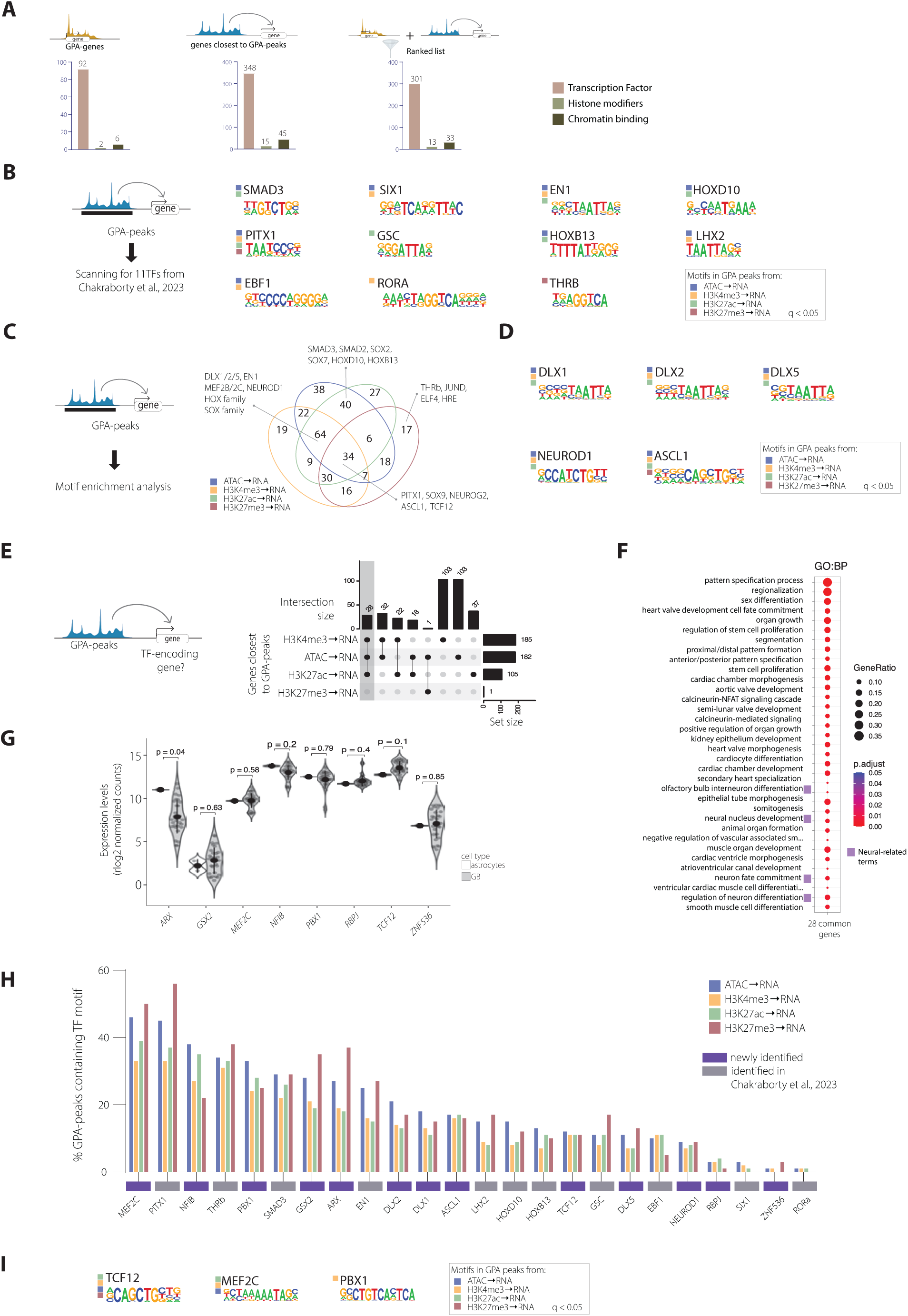
Identification of transcription factors linked to GB-relevant regulatory networks. **(A)** Panther annotation of GPA-genes (left), genes closest to GPA-peaks (middle) and ranked-list genes (right) as transcription factors (TFs), histone modifiers and chromatin binding proteins. **(B)** Scanning for the motifs of 11 key TFs identified in Chakraborty et al., 2023 [7] within the GPA-peaks (left), and significantly enriched motifs for the 11 TFs scanned within GPA-peaks (q<0.05) (right). **(C)** Motif enrichment analysis of the GPA-peaks (left) identifies additional TFs relevant for GB (right), including various TFs associated to synaptic functions. **(D)** Motifs significantly enriched within GPA-peaks for those TFs related to GABA signaling as in (C) (q<0.05). **(E)** TF-encoding genes closest to GPA-peaks (left) were intersected across the four lists (right), identifying 28 TF-encoding genes whose promoters are characterized by chromatin accessibility and the presence of active marks. **(F)** GO analysis of the gene set identified in (E) reveals enrichment for GO terms related to neural function. **(G)** Gene expression levels of the 8TF-encoding genes belonging to the neural-related GO terms identified in (F). **(H)** Percentage of the GPA-peaks containing motifs for the TFs selected from (B), (C) and (G). **(I)** Motifs significantly enriched within GPA-peaks for TFs related to neural functions as in (C) (q<0.05).

Next, we wondered whether MOBILE could also capture the key 11 TFs identified in our previous study for their relevance in the rewiring of transcriptional programs in GB [11]. For this purpose, we scanned the GPA-peaks for the motifs of these 11 keys TFs and determined the motif occurrence within the peaks (Fig. 4B, left). The motifs of all 11 TFs were significantly enriched within the GPA-peaks (Fig. 4B, right). Importantly, PITX1, THRB and SMAD3 stood out at the top when ranking the TFs based on the percentage of GPA-peaks containing motifs for each TF of interest (Fig. 4H). These results align with our previous study, where SMAD3 and PITX1 were identified as the major TFs involved in a GRN underlying glutamatergic neuron-to-glioma synaptic communication [11]. In addition to this, motif enrichment analysis of GPA-peaks revealed a significant enrichment for PITX1, SOX9, NEUROG2, ASCL1 and TCF12, among others, across all four association lists (Fig. 4C,D and Supplementary Data 2). SOX9 expression is generally elevated in GB tumour biopsies, correlating with poor prognosis [64]. NEUROG2 functions as a proneural gene and is critical for specifying glutamatergic neural identity in the dorsal telencephalon [65]. The decision between glutamatergic and GABAergic fates in telencephalic progenitors is governed by a core transcriptional network, in which the regional expression of Neurog2 and Ascl1 determines neuronal subtype identity, with Neurog2 promoting glutamatergic differentiation and Ascl1 promoting GABAergic fate [65]. Other motifs significantly enriched at GPA-peaks are those for the DLX, HOX and SOX TF families, as well as EN1, MEF2B/C, NEUROD1 (Fig. 4C, Supplementary Data 2). DLX1, DLX2, and DLX5 are essential TFs in GABAergic signalling, particularly in the development, function and maintenance of inhibitory GABAergic interneurons [66,67]. DLX1 and DLX2 drive the synthesis of GABA by regulating *GAD1/2*, encoding critical enzymes for GABA production, as well as promoting synaptogenesis and dendritogenesis to support inhibitory synaptic networks [66]. GABA-induced expression of *NeuroD1* in glial progenitors is associated with increased histone H4 acetylation, suggesting that GABA not only promotes neuronal differentiation through NeuroD1 but also integrates epigenetic mechanisms, such as histone acetylation, to regulate progenitor cell fate [68]. Noteworthy, the percentage of GPA-peaks containing motifs for NEUROD1, ASCL1 and DLX was substantial, though lower than the observed for SMAD3 and PITX1 (Fig. 4H).

To further explore this dataset, we examined the genes closest to GPA-peaks that encode for TFs. Out of the total, 28 genes encoding TFs were common to the associations involving chromatin accessibility and active histone marks (Fig. 4E). GO analysis revealed significant enrichment for terms related to interneuron differentiation, neuron fate commitment and neuron differentiation (Fig. 4F). Out of the 28 TF-encoding genes, 8 are involved in these neuronal processes, including *ARX, GSX2* and *MEF2C,* and all are expressed in GB (Fig. 4G). ARX and GSX2 have been linked to the differentiation of GABAergic interneurons [69–71]. Mutations in *ARX* results in neural defects linked to GABAergic neurons [69], and in mouse *Arx* expression is regulated by *Dlx* genes [72] . In particular, Arx is a direct target of Dlx2, and it contributes to the migration of GABAergic interneurons [72]. Moreover, Dlx1/2 and Gsx2 are essential for promoting and maintaining the expression of *GAD1*, central for GABAergic neurons [73,74]. Additionally, the percentage of GPA-peaks containing motifs for ARX and GSX2 is similar to the observed for SMAD3, though lower than for PITX1, while MEF2C shows the highest occurrence (Fig. 4H). In the healthy brain, MEF2C is essential for neuronal development, synaptic plasticity, and cognitive function, highlighting its role in neurodevelopment and neuronal circuit maintenance [75,76]. In brain metastasis, *MEF2C* expression and nuclear translocation have been associated with disease severity and proliferative activity [77]. Notably, ARX has been reported to control genes involved in cell specification like *Mef2c,* among other genes linked to cell cycle and migration [70]. In addition to MEF2C, the motifs for TCF12 and PBX1 are also significantly enriched within GPA-peaks (Fig. 4I). TCF12 and NeuroD1 cooperate to establish active chromatin states, particularly at loci governing neural migration during cortical development [78]. Noteworthy, PBX1 and PAX6 are key components in a transcriptional network that directs adult-born neural progenitor cells towards the acquisition of a GABAergic phenotype [79].

Altogether, our integration of multi-omics data, obtained from 15 patient-derived GB lines alongside normal astrocytes, uncovered novel GRNs relevant for GB pathogenesis and identified the TFs potentially operating through these networks. GABA-related signalling emerged as a previously unrecognized driver of GB pathogenesis, and TFs such as ARX, GSX2 and DLXs were identified as relevant regulators in this context. Following the identification of GABA-related genes by MOBILE, we revisited the topological and epigenetic landscape of their loci, as mapped in our previous study [11]. The loci of genes such as *GAD1, ASCL1, NEUROG2, MEF2C, ARX or DLX1/2,* among others, undergo extensive changes regarding histone modifications, chromatin accessibility, and chromatin loops (Supplementary Fig. 7). These findings underscore an epigenetic reprogramming that likely drives alterations in GABA-related pathways, reinforcing the role of GABA signalling in GB progression.

### Effect of glutamatergic and GABAergic neural activity on GB cell proliferation

Glutamatergic AMPA receptor-dependent synapses between neurons and glioma cells have been observed in paediatric and adult high-grade gliomas [7,8]. Noteworthy, these glutamatergic neuron-to-glioma synapses have been identified as key drivers of glioma progression, including GB. However, the contribution of GABAergic signalling to GB remains a topic to be investigated. In diffuse midline gliomas (DMG), a type of pediatric high-grade glioma, tumour-promoting GABAergic neuron-to-glioma synapses have been recently identified [47]. Since our findings highlighted the involvement of GABA-related networks in GB, we sought to functionally test this using *in vitro* co-cultures of GB cells and human embryonic stem cell (hESC)-derived neurons, both glutamatergic and GABAergic, specifically aiming to address how neuron-GB interactions influence GB cell proliferation.

For this purpose, we first differentiated hESCs into regionalised neural progenitors (i.e., dFB progenitors are dorsal telencephalon and MGE cultures are ventral telencephalon based on the expression of neural markers FOXG1, PAX6, NKX2.1 and ISL1, Supplementary Fig. 8A), that further matured into either glutamatergic neurons or GABAergic interneurons over a total of 42 days (Fig. 5A, B), timepoint at which they display expression of the neural marker beta III-tubulin (Fig. 5C). GFP-tagged GB cells (i.e., U251-GFP) were seeded directly onto the neuron layer in a co-culture set-up at day 42, and proliferation of GB cells was monitored over 48 h by live-cell imaging (Fig. 5D-I). Compared to standalone monocultures, GB cells exhibited increased proliferation in co-culture not only with glutamatergic neurons -as expected based on previous reports-, but also with GABAergic interneurons (Fig. 5D-E). These results indicate that while both neuron types seem to influence GB proliferation, glutamatergic signalling might play a more prominent role than GABAergic input. Next, we assessed the impact of neuron-specific pharmacological treatments on GB cell proliferation in the co-culture assays. In co-culture with glutamatergic neurons, treatment with NBQX, a competitive antagonist of AMPA receptors, significantly reduced the proliferation of GB cells (Fig. 5F, G). This finding aligns with the reported effect of glutamatergic neuron-to-glioma synapses on promoting GB progression [7,8]. This effect is specifically observed in the co-culture setting, as NBQX treatment did not affect the growth of GB cells in monoculture (Supplementary Fig. 8B). We then investigated whether inhibition of GABA receptors (GABAR) could similarly influence GB cell proliferation. Based on the presence of GABA_A_R-dependent GABAergic neuron-to-glioma synapses in DMG [47], and the broad expression of GABA_A_R genes in GB cells (Supplementary Fig. 8C and [11], we pharmacologically inhibited GABA_A_R using picrotoxin, a non-competitive antagonist. However, picrotoxin had no effect on the proliferation of GB cells in co-culture with GABAergic interneurons (Fig. 5H,I), suggesting that GABAergic input does not contribute through classical synaptic mechanisms.

**Figure 5.**
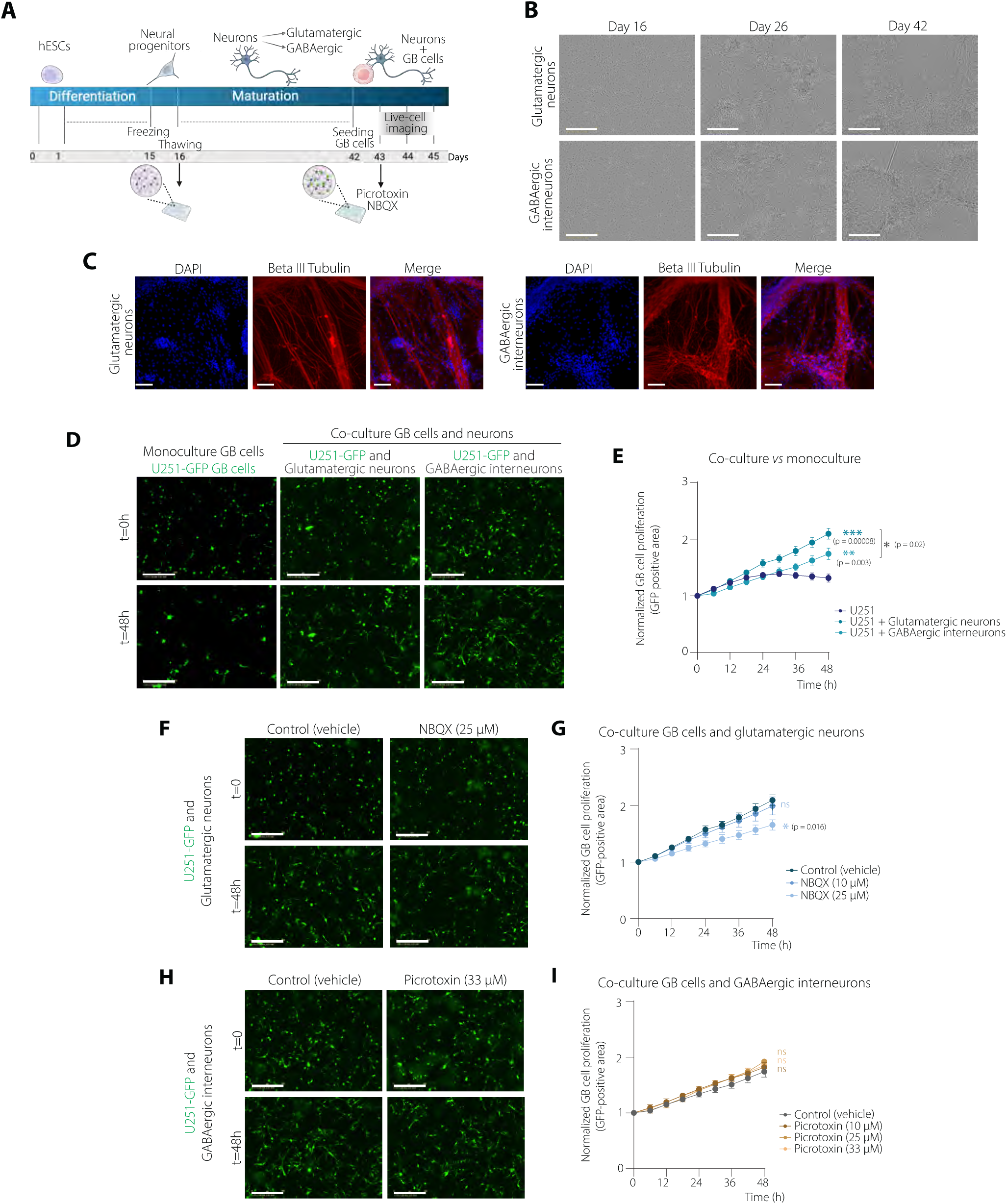
Effect of glutamatergic and GABAergic neural activity on GB cell proliferation. **(A)** Schematic representation of the experimental design. hESC-derived neurons (glutamatergic neurons or GABAergic interneurons) were co-cultured with GFP-tagged U251 GB cells and subjected to different drug treatment combinations. **(B)** Representative brightfield images of glutamatergic neurons (top panels) and GABAergic interneurons (bottom panels) over time (days 17, 26, and 42). Scalebar=400 µm. **(C)** Immunostaining of glutamatergic neurons or GABAergic interneurons with a beta III tubulin antibody on day 42. Scalebar=100 µm. **(D)** Microscopy images showing U251-GFP cell proliferation for U251-GFP cells in mono-culture (left), or in co-culture with glutamatergic neurons (middle) or GABAergic interneurons (right), at t=0 h and t=48 h. Scalebar=400 µm. **(E)** Proliferation of GB U251-GFP cells over 48 h in co-cultures with glutamatergic neurons or GABAergic interneurons compared to U251-GFP cells cultured alone. Data is represented as GFP-positive area over time and normalized to t=0 h, and presented as mean ± SEM (n=6). Statistical significance was assessed by unpaired t-test with Welch’s correction (*p<0.05, **p<0.005, ***p<0.0005). **(F-I)**: Representative fluorescence images and corresponding proliferation curves for U251 GFP-positive cells in co-culture with glutamatergic neurons (F, G) or GABAergic interneurons (H, I), following the indicated treatments [NBQX (10 µM, 25 µM), picrotoxin (10 µM, 25 µM, 33 µM)] compared to vehicle-control. GB cell proliferation was determined as GFP-positive area, normalized to t=0h and displayed as mean ± SEM. Statistical significance was assessed by unpaired t test with Welch’s correction (*p<0.05, **p<0.005, ***p<0.0005). Scale bars=100 µm.

Altogether, our data show that both glutamatergic and GABAergic neurons can enhance GB cell proliferation *in vitro* and may contribute to a pro-tumorigenic environment in GB. Noteworthy, these findings also suggest that while both neuron types can promote GB cell proliferation, their modes of action and underlying mechanisms may differ. The lack of response to GABA_A_ receptor inhibition suggest that GABAergic signalling may influence GB cell proliferation through pathways distinct from classical synaptic transmission. GABAergic modulation of GB cells could instead involve non-canonical pathways or indirect mechanisms, such as metabolic or paracrine interactions.

## DISCUSSION

Integrating multi-omics data with the ML-based MOBILE pipeline, this study uncovers novel gene regulatory networks (GRNs), and pathways involved in glioblastoma (GB) pathogenesis. Among the most notable findings, GABAergic signalling emerged as a previously unrecognized driver of GB malignancy. MOBILE consistently identified GABA-related networks and transcription factors (TFs) such as ARX, GSX2, and DLX family members, known for their key roles in the development and function of GABAergic interneurons. These results expand our understanding of GB biology, highlighting that GABAergic signalling, like the well-established glutamatergic pathways, may also contribute to glioblastoma progression.

Neuron-glioma interactions have emerged as a critical area of study, shedding light on the intricate ways in which the nervous system influences cancer progression. Recent studies have highlighted that GB cells can integrate into neural circuits, receiving synaptic input and leveraging neuronal signals to promote tumour growth and brain invasion [7,8,10,80]. This occurs through different mechanisms including paracrine signalling [9,38,39] and glutamatergic neuron-to-glioma synapses [7,8]. Our group recently showed that changes in 3D genome organization and the epigenetic landscape in GB orchestrate gene expression programs underlying neuron-to-glioma synaptic communication, and identified the major TFs and regulatory networks that mediate this neuron-glioma crosstalk [11]. The current study not only reinforces the relevance of these pathways, but more importantly reveals additional regulatory networks that point to GABAergic signalling as an important contributor to GB.

GABAergic synapses have been recently identified in diffuse midline gliomas (DMG), while only minimal depolarizing GABAergic currents were found in hemispheric high-grade gliomas, such as IDH-wildtype glioblastoma [47]. Noteworthy, diffuse hemispheric gliomas were shown to adopt GABAergic interneuron-like transcriptional profiles [46], and a defined group of interneuron progenitors can give rise to certain brain tumours [81]. GABAergic signalling has also been linked to other cancers. Upregulation of *GABRB3*, for instance, has been associated with the occurrence of brain metastases originating from diverse cancers, including prostate and breast cancer [82]. Breast-to-brain metastatic tissue and cells display a GABAergic phenotype similar to that of neuronal cells [83]. Adopting this GABAergic phenotype could facilitate tumour adaptation and progression through different pathways, such as potentially PI3K/AKT which becomes activated upon stimulation of GABRB3 by GABA, thus regulating cell survival, anti-apoptosis and proliferative pathways [82].

Notably, GABA is not only used as a signalling molecule but also as a nutrient. In medulloblastoma, GABA is utilized as a metabolic substrate and metabolized to support tumour growth [84]. Vigabatrin, an irreversible inhibitor of GABA transaminase, reduces brain metastases by blocking GABA catabolism, thereby depriving tumour cells of key metabolic intermediates despite elevated GABA levels [63]. One could speculate that GB may co-opt GABAergic signalling not only for energy metabolism using glutamine, via increased *GAD1* expression, but also for oncogenic neuron-to-glioma communication. In this context, our *in vitro* co-culture experiments provided further insights into how neurons influence GB cell behaviour. In both the glutamatergic and the GABAergic setting, GB cell proliferation increased compared to monoculture, indicating that neuronal input -regardless of neurotransmitter identity-supports tumour growth, though seemingly through different mechanisms. Pharmacological blockade of AMPA receptors with NBQX reduced GB proliferation in co-culture with glutamatergic neurons, consistent with the tumour-promoting role of glutamatergic synaptic input. However, picrotoxin, a GABA_A_ receptor antagonist, had no effect on GB proliferation in co-culture with GABAergic neurons, suggesting that GABAergic input may act through non-synaptic pathways, potentially including metabolic or paracrine mechanisms. These findings point to GABA metabolism as a potential therapeutic vulnerability in GB, highlighting the translational potential of GABA-targeting drugs, such as vigabatrin, to disrupt metabolic adaptations that may sustain tumour growth in the brain.

In addition to GABAergic signalling, our study reinforces the relevance of other GRNs linked to neurotransmission, including glutamatergic signalling. Consistent with prior studies, MOBILE captured key pathways, such as those related to synapse organization, axon guidance and glutamatergic synapses, that support glioblastoma’s ability to integrate into neural circuits [11,80,85]. NLGN3, which was reported to promote high-grade glioma proliferation [38,39], was identified by MOBILE as relevant for GABA-related networks. The fact that NLGN3 localizes at both excitatory and inhibitory synapses [59,60], urges further studies to address the mechanistic role of GABAergic signalling in GB pathogenesis. These findings also highlight the importance of understanding these mechanisms to design efficient strategies to target neuron-glioma interactions as a therapeutic strategy in glioblastoma.

Beyond neurotransmission, MOBILE identified GRNs linked to ECM remodelling and immune evasion as relevant for GB progression. The enrichment of immune-related pathways, particularly those involving MHC components, suggests that GB could employ strategies to evade immune detection by downregulating MHC class I expression. In glioblastomas, altered MHC class I expression can modify NK cell function, as lower MHC I levels may decrease NK cell inhibition, thereby facilitating their activation [45]. This also aligns with the immune evasion strategies seen in other cancers, where downregulation of MHC class I molecules helps tumour cells escape detection [86]. Moreover, HLA-B/C expression correlates with immune evasion and poor prognosis in colorectal cancer [49]. Gaining insight into these mechanisms can aid in developing effective immunotherapies for GB. Additionally, we identified networks related to extracellular matrix (ECM), emphasizing the role of ECM remodelling in GB. The downregulation of LOXL4 observed in our patient-derived GB lines relative to normal astrocytes, despite its upregulation in TCGA data [87], suggests that tumour microenvironment inputs may influence its expression. Given the role of LOXL4 in tissue stiffening [88] and inhibition of Ras/ERK signalling [51], its differential expression across datasets warrants further investigation into its context-dependent functions. MMP14 (matrix metalloproteinase 14) facilitates ECM remodelling, promoting tumour invasiveness and metastasis [57], while the fibrillar collagen COL11A1 is associated with chemoresistance and poor outcomes in epithelial cancers like ovarian and pancreatic cancer [58]. Our analysis also detected networks involving *FN1* (fibronectin 1), supporting cell adhesion and migration, both essential for metastasis [54]; *TGFBI* (transforming growth factor-beta-induced protein), which influences tumour progression by regulating epithelial-mesenchymal transition and immune evasion [55]; as well as *SPARCL1* (SPARC-like protein 1), often recognized as a suppressor of metastasis given its role in cell-cell and cell-matrix interactions [56].

Altogether these results highlight the capability of MOBILE to integrate high-dimensional multi-omics data and identify novel epigenetically regulated genes pivotal in oncogenic processes. This study, while confirming the relevance of glutamatergic-related networks, also reveals the importance of novel GABA-linked GRNs. GABAergic signalling has thus been identified as a previously unrecognized driver of GB malignancy, through the discovery of GABA-related networks and relevant TFs that regulate GABAergic pathways. This finding is further supported by the *in vitro* co-culture assays showing that GABAergic input promotes the proliferation of GB cells, potentially through metabolic or paracrine mechanisms rather than synaptic transmission. Our findings call for further investigations into the role of GABA in GB, particularly to elucidate potential metabolic or paracrine pathways, or to determine whether neuron-to-glioma GABAergic synapses might exist but remain undetected. In either case, these insights will help inform the development of strategies to target neuron-to-glioma interactions in such a difficult-to-treat cancer as glioblastoma.

## MATERIAL AND METHODS

### Source of Multi-Omics Datasets and Mapping

Multi-omics datasets for analyses of Gene Regulatory Networks (GRNs) were previously generated by our group [11]. These data include RNA-seq, ATAC-seq, and ChIP-seq for histone modifications H3K27ac, H3K27me3 and H3K4me3 from 15 patient-derived glioblastoma cell lines alongside normal human astrocytes. Fastqfiles were quality-checked with FastQC (0.11.8). RNA-seq raw reads were mapped to the human genome (GRCh38/hg38) using STAR (2.7.6b), and genes with a minimum row sum of 10 reads were kept for further analysis. ChIP-seq raw reads were mapped to the human genome (GRCh38/hg38) using bowtie2 (2.4.1) (--threads 4 --very-sensitive). The processing of ATAC-seq raw data included trimming with cutadapt (2.10) to remove the Nextera adaptor sequence CTGTCTCTTATACACATCT and improve mapping quality, mapping to the human genome (GRCh38/hg38) using bowtie2 (--k 2, --threads 8, -- local, --maxins 2000), removing duplicates with MarkDuplicates from Picard toolbox (2.27.5) and filtering with samtools (1.12) to keep high-quality and uniquely aligned read pairs. To prepare these datasets for MOBILE [37] analysis, BAM files were processed to generate pseudo-replicates using SAMtools v1.15.1 [89], yielding pseudo-replicates ∼20-50 million read pairs each.

### Peak Calling and Consensus Peaks

Peak calling for ChIP-seq and ATAC-seq was performed by MACS3 (3.0.0b1) (options: -f BAMPE --broad --nomodel --shift -100 --extsize 200 -B -g hs -q 0.05). Consensus peaks were generated for each sample to identify non-redundant peak sets. BAM files were processed in R (4.4.0) using the GenomicRanges package v1.56.2 [90]. Peaks were retained if they overlapped with peaks from each pseudo-replicate by more than 50%. Read counts within consensus peaks were quantified using the Rsubread package v2.18.0 [91], producing peak-level counts that were subsequently normalized using the DESeq2 package (v1.12.2, Bioconductor v3.19.1) to obtain rlog2 normalized counts per sample.

### Feature Selection and Data Filtering

To identify molecular features associated with cellular phenotypes and pathways, the MOBILE (Multi-Omics Binary Integration via Lasso Ensembles) framework was applied to the rlog2 normalized counts [37]. Prior to analysis, a raw variance filter was applied to each assay to retain only the top 5% of highly variable features.

### Multi-Omics Data Integration with MOBILE

#### Lasso Regression Analysis

To assess epigenetic influences on gene expression, we used the Lasso module within MOBILE to compute matrix *β* in the equation *Y*=*β*⋅*X*+*δ*, where Y and X represent RNA-seq and ATAC/ChIP-seq matrices, respectively. Four epigenetic-transcriptomic relationships were analysed: ATAC-seq→RNA-seq, H3K27ac ChIP-seq→RNA-seq, H3K27me3 ChIP-seq→RNA-seq, and H3K4me3 ChIP-seq→RNA-seq, yielding four corresponding coefficient matrices. To identify robust predictors, Lasso regression is independently run 10,000 times. Coefficients that appear in at least half of these 10,000 iterations (i.e., >5,000 iterations) are considered stable and retained. The resulting matrix of these consistently selected coefficients is referred to as the Robust Lasso Coefficient Matrix (RLCM), which is used for further downstream analysis.

#### Glioblastoma-specific Analysis using LOGO

To identify glioblastoma-specific associations, we utilized MOBILE’s LOGO (Leave-One-Group-Out) module. LOGO analyses were performed by excluding the data from the normal human astrocytes and comparing the results to the complete dataset (FULL-data), generating coefficient matrices that reflect glioblastoma-specific (GB-specific) associations. This analysis was conducted for each of the four epigenetic-transcriptomic pairs defined above.

### Construction of the Integrated Associations Network (IAN)

#### Network Generation and Output Description

Following Lasso and LOGO analyses, the resulting coefficient matrices were merged to construct a gene-level network termed the Integrated Associations Network (IAN). In this network, nodes correspond to individual genes, while edges represent inferred associations between genes and epigenetic or transcriptomic features. The strength and directionality of these associations are captured by edge weights, which are based on the corresponding Lasso coefficients. The IAN provides a systems-level view of gene-feature interactions, highlighting clusters of genes with strong shared interactions (i.e., high coefficient values). These clusters may reflect co-regulated genes or components of common biological pathways (see [37] for further details). For each pairwise comparison, two types of outputs were generated:

1. *Associations list:* A list of epigenetic-transcriptomic associations i.e., association between a ChIP/ATAC peak and the expression of a gene, hereby named Gene-Peak Associations (GPA). This list contains the RNA-seq gene name (i.e, GPA-gene), the gene name closest to the associated ChIP/ATAC peak (i.e., gene closest to GPA-peak), peak coordinates, and coefficient values for both FULL-data and LOGO models.
2. *Ranked list:* A ranked gene list, where each row contains a gene (network node) and the weighted sum of all edge weights for that gene, representing the magnitudes of the Lasso coefficients. Any association (lasso coefficient) with a magnitude smaller than 0.01 was excluded from consideration.

### Pathway Enrichment and Network Visualization

#### Pathway and Gene Ontology Enrichment

Pathway enrichment analysis was conducted on each ranked gene list using Gene Set Enrichment Analysis (GSEA) (4.3.3) [92], with the C5 Gene Ontology Sets BP, CC and MF (v2023.1.Hs) as the gene set database and the Enrichment Map Visualization tool of GSEA together with Cytoscape (v3.10.2), and applying a significance threshold of p<0.05 and FDR<0.1. Gene Ontology (GO) enrichment was performed using the clusterProfiler package v4.12.6, applying a significance threshold of p<0.01. Tree plots for GO visualization were created using the ggtree package v3.12.0.

#### Network Visualization

Integrated networks were visualized in Cytoscape v3.10.1 [93]. Clustering of differentially expressed genes was performed in R using the pheatmap package v1.0.12, and gene intersections among ranked lists were visualized using the UpSet command from the ComplexHeatmap package v2.20.0. Violin plots of RNA-seq data (rlog2 normalized counts) were generated using ggplot2 v3.5.1, ggpubr v0.6.0, and rstatix v0.7.2.

### Transcription Factor Analysis

#### Identification of Relevant Transcription Factors

GPA-genes and genes close to GPA-peaks, retrieved from the associations lists, alongside genes in the ranked lists were assessed for annotations as transcription factors (TFs), histone modifiers or chromatin binding using the Panther Database [94].

#### Transcription Factor Binding (TFBS) Motif Analysis

TFBS motifs within GPA-peaks were identified using HOMER (v4.11.1) findMotifsGenome.pl tool. GPA-peaks were also used to scan for motifs of particular TFs using their positional weight matrices (PWMs), with default parameters (-size 200, -mask). The selected TFs included EBF1, EN1, GSC, HOXB13, HOXD10, LHX2, PITX1, RORA, SIX1, SMAD3, THRB, ARX, GSX2, MEF2C, NFIB, PBX1, RBPJ, TCF12, and ZNF536.

### Cell Culture

All cultures were maintained in a cell incubator at 37°C in a humidified atmosphere (95% humidity) with 5% CO_2_.

#### Glioblastoma cells

U251 glioblastoma cells (Sigma-Aldrich, #09063001, authenticated by STR profiling) were grown in EMEM (EBSS) supplemented with 2 mM Glutamine, 1% NEAA (Non-Essential Amino Acids), 1 mM Sodium Pyruvate, 10% FBS (Fetal Bovine Serum) and 1% penicillin/streptomycin (all from Gibco). The GFP-labelled U251 line was established by lentiviral integration of the GFP reporter gene.

#### hESC-derived glutamatergic neurons and GABAergic interneurons

RC17 (Roslin Cells, hPSCReg: RCe021-A (RRID:CVCL_L206)) hESCs were differentiated for 16 days into medial ganglionic eminence (MGE) GABAergic interneuron progenitors and dorsal forebrain (dFB) glutamatergic progenitors. Directed differentiation of MGE GABAergic progenitors was obtained as previously described [95]. At day 0 of differentiation, cells were seeded at 25,000 cells/cm^2^ in N2 medium consisting of N2 supplement (1:100 cat. 17502048 Thermo Fisher Scientific), DMEM/F12 (0.5x cat. 11320033 Thermo Fisher Scientific), Neuromedium (0.5x cat. 130-093-570 Miltenyi Biotec) and glutamax (1:100 cat. 35050061 Gibco) supplemented with ROCK inhibitor (10µM 10 μM Y-27632, Cat. 130-106-538 Miltenyi Biotec) for the first 48h and growth factors as described next. GABAergic progenitors were differentiated in the presence of SB431542 (10 µM, Miltenyi Biotec Cat. 130-106-543) day 0-9, Noggin (100 ng/ml, Miltenyi Biotec Cat. 130-108-982) day 0-9 and XAV939 (3 µM, StemCell Technologies Cat. 72674) day 0-9 and SHH-C24II (200 ng/ml, Miltenyi Biotec Cat. 130-095-730) day 3-14. Differentiation of dFB glutamatergic progenitors were obtained by exposure to Noggin (100 ng/ml) and SB431542 (10 µM) from day 0-4 for both. After 16 days of differentiation the neural progenitors were cryopreserved at 4-6 million cells per vial in Cryostor CS10 (StemCell Technologies Cat. 07930). Thawed progenitors were seeded onto 96-well plates (Sarstedt, Cat. 83.3924) coated with 2 μg/cm^2^ of human recombinant laminin 521 LN (Biolamina, Cat. LN521-05) in DPBS+Ca+Mg (Gibco, 14040117), in NB21 media, including neuromedium (Miltenyi, Cat. 130093570), neurobrew-21 (Miltenyi, Cat. 130097263), 10 U/ml penicillin/streptomycin (Gibco,

Cat. 15140122), 2 mM glutamax (Gibco, Cat. 35050061) and supplemented with factors: 0.2 mM ascorbic acid (Sigma, Cat. A4403), 20 ng/ml BDNF (Miltenyi, Cat. 130-093-811), 500 μM cAMP (Sigma, Cat. D0627), 1 μM DAPT (Miltenyi, Cat. 2634/10) and 10 μM Rock inhibitor (Miltenyi, Cat. 130-106-538). The cells were counted using Countess II FL Automated Cell Counter (Thermofisher) and seeded at a density of 270,000 cells/cm^2^. Neurons were maintained by full media change (NB21 media with supplements) every second day for the first 7 days and thereafter 75% media change for the rest of the maturation time.

#### Contact co-culture of hESC-derived neurons and GB cells

To establish the contact co-culture assay, 5000 U251 GFP-positive GB cells were seeded onto day 42 hESC-derived neurons and allowed to settle. GB monocultures and co-cultures were then treated with either the AMPA receptor antagonist NBQX (10 or 25 µM 2,3-Dioxo-6-nitro-1,2,3,4-tetrahydrobenzo[*f*] quinoxaline-7-sulfonamide, Tocris Bioscience #1044) or the GABA_A_ receptor antagonist picrotoxin (10, 25 or 33 µM, Tocris Bioscience #1128). Co-cultures were maintained as described above and monitored by live-cell imaging using the IncuCyte S3 Live-Cell Analysis instrument (Sartorius) to assess cell proliferation over a 48 h period. Proliferation of the GFP-positive GB cells was determined by measuring GFP-positive area every 6 h using the Incucyte Base Analysis Software (Sartorius). Significant differences in cell proliferation were assessed using unpaired t-test with Welch correction at t =48h (*p <0.05, **p <0.005, ***p < 0.0005).

### Immunolabelling

#### Immunofluorescence on neural progenitors

Immunolabelling of neural progenitors was performed as previously described [96]. Briefly, cells were seeded onto coated 96-well plates, using the coating and culture conditions as described above. For immunofluorescence, cells were washed once in 1x PBS and fixed in 4% PFA (Thermo Fischer Scientific, Cat. 28908) for 15min at room temperature (RT), followed by three additional washes in 1x PBS, and incubation in blocking buffer containing 5% donkey serum (VWR, Cat. S2170-100) in 1x PBS, 0.1% TritonX-100 (Sigma Aldrich, Cat. T8787) and 0.02% NaN_3_ (Sigma Aldrich, Cat. S2002-5G) for 1-3h at RT. Incubation with primary antibodies diluted in blocking buffer was performed overnight at 4°C on a shaker, as follows: FOXG1 (1:500, rabbit, Abcam ab18259), PAX6 (1:1000, mouse, Sigma-Merck, Cat. AMAb91372), NKX2-1 (1:100, rabbit, Abcam ab133737) and ISL-1 (1:50, mouse, DSHB, Cat. 39.4D5). After three washes in 1x PBS, cells were incubated for 2h at RT with the corresponding secondary antibodies (Donkey anti-rabbit-AF647, 1:200, Jackson ImmunoResearch Cat. 711-605-152; or Donkey anti-mouse-AF488, 1:200, Jackson ImmunoResearch Cat. 715-545-150), diluted in blocking buffer containing DAPI (1:1000, Thermo Fisher Scientific Cat. D3571). After three final washes with 1x PBS, cells were stored in 0.02% NaN_3_ in 1x PBS at 4°C and protected from light. Images were acquired on the Leica AF6000 fluorescence microscope and image analysis was performed using the Fiji software.

#### Immunofluorescence on hESCs-derived neurons

For immunofluorescence purposes, glutamatergic and GABAergic progenitors were thawed and seeded onto coated 18-well chambered glass coverslips (IBIDI, Cat. 81817) at a density of 270,000 cells/cm^2^, using the coating and culture conditions as described above. On day 42, cells were washed with 1X PBS (pH 7.4) and fixed in 4% PFA (Thermo Scientific, Cat. 28906) for 15 min at RT, followed by 3x 5min 1X PBS washes. Permeabilization was performed with 0.2% Triton-X100 (Sigma Aldrich, Cat. T8787) in PBS for 5min on ice, followed by 3x 5min washes in 1X PBS. Blocking was done in 10% FBS (Life technologies, #10500064) in PBS for 30 min at RT in a dark humid chamber. Cells were then incubated overnight at 4°C with the primary antibody (beta III tubulin, 1:1000, Abcam ab18207) diluted in blocking solution, thereafter washed 3x 5min with PBS, and incubated with secondary antibody (Goat anti-Rabbit Alexa Fluor 594, 1:500, Thermo Scientific, Cat. A32740) for 2h at RT, followed by 3x 5min PBS washes. Nuclei were counterstained with DAPI (1:1000, Cat. ab285390) for 5 min, washed again, and mounted using IBIDI mounting media (Cat. 50011). Slides were stored at 4 °C, and image acquisition was performed on a Leica Thunder inverted widefield microscope.

## DATA AVAILABILITY

The raw RNA-seq, ChIP-seq and ATAC-seq datasets, from both patient-derived glioblastoma lines and normal human astrocytes, were generated by [11] and available at GEO (Gene Expression Omnibus) under the accession number GSE217349 (https://www.ncbi. nlm.nih.gov/geo/query/acc.cgi?acc=GSE217349).

## SUPPLEMENTARY DATA

Supplementary Data are available online.

## AUTHOR CONTRIBUTIONS

I.N. performed all the computational analyses, interpreted data, prepared figures and drafted the original manuscript. S.D. and C.C. performed neural cultures and co-culture assays, and analysed and interpreted the corresponding data. A.H.N. differentiated hESCs into neural progenitors (glutamatergic and GABAergic) and characterized the progenitors by immunolabeling under supervision of A.K. A.H. co-supervised some analysis, and contributed to data interpretation and manuscript editing. C.E. co-supervised the MOBILE-based computational analyses and contributed to manuscript editing. S.R. designed and supervised the study, secured funding, analysed and interpreted the data, and wrote the final manuscript with input from the other authors.

## ACKNOWLEDGEMENTS

The computations were enabled by resources in projects snic2020-15–157, snic2021-22-504 and naiss2023-22-55 provided by the Swedish National Infrastructure for Computing (SNIC) and the National Academic Infrastructure for Supercomputing in Sweden (NAISS) at UPPMAX, funded by the Swedish Research Council through grant agreements no. 2018-05973 and no. 2022-06725, respectively, alongside resources from High Performance Computing Center North (HPC2N) hpc2nstor2023-051 and hpc2n2024-025. We also acknowledge the Biochemical Imaging Centre at Umeå University and the National Microscopy Infrastructure, NMI (VR-RFI 2019-00217) for providing assistance in microscopy.

