## Supplementary Information for "Gene regulatory networks linked to GABA signalling emerge as relevant for glioblastoma pathogenesis"

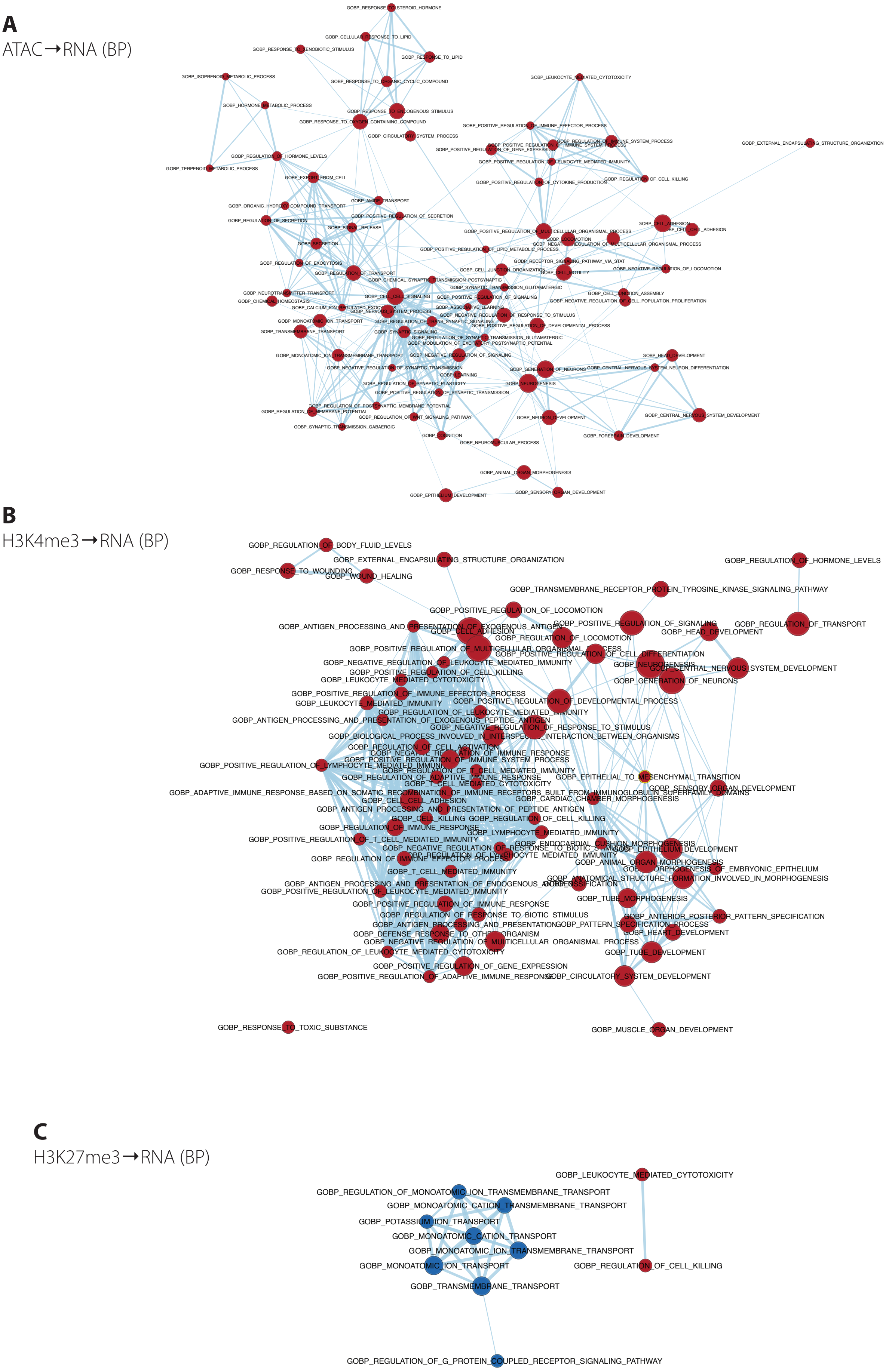


**Supplementary Figure 1. Gene regulatory networks identified by MOBILE.** Pathways identified as significant for GB, with a p-value cutoff of 0.05, for ATAC→RNA-seq **(A)**, H3K4me3→RNA-seq **(B)**, and H3K27me3→RNA-seq **(C)**, for biological processes (BP).

**A**

ATAC→RNA (CC)

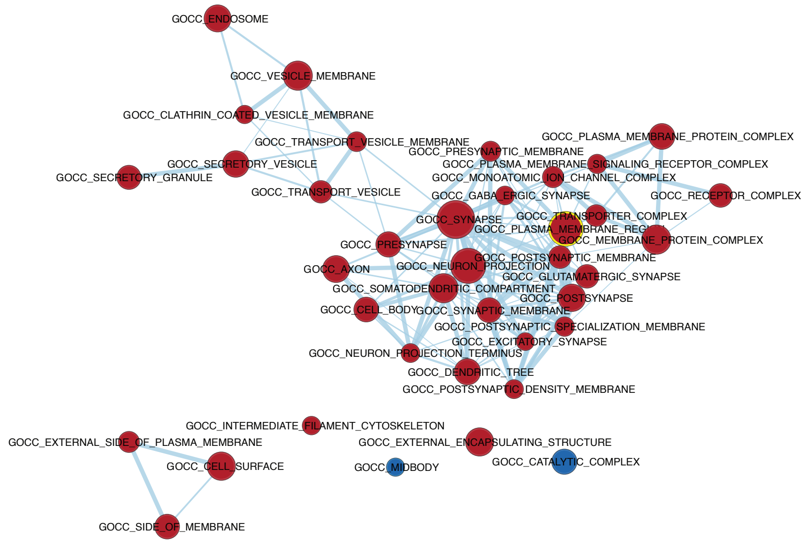


**B**

H3K4me3→RNA (CC)

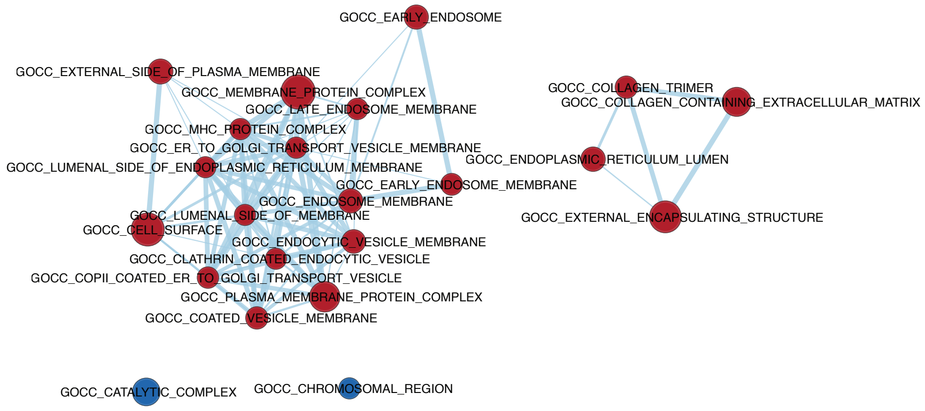


**C**

H3K27ac→RNA (CC)

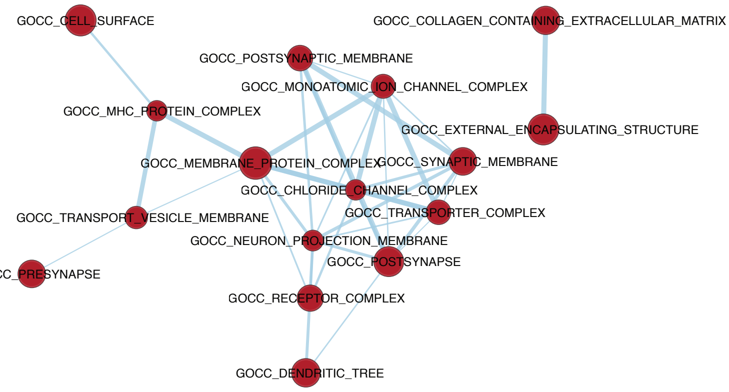


**Supplementary Figure 2. Gene regulatory networks identified by MOBILE.** Pathways identified as significant for GB, with a p-value cutoff of 0.05, for ATAC→RNA-seq **(A)**, H3K4me3→RNA-seq **(B)**, and H3K27ac→RNA-seq **(C)**, for cellular component (CC).

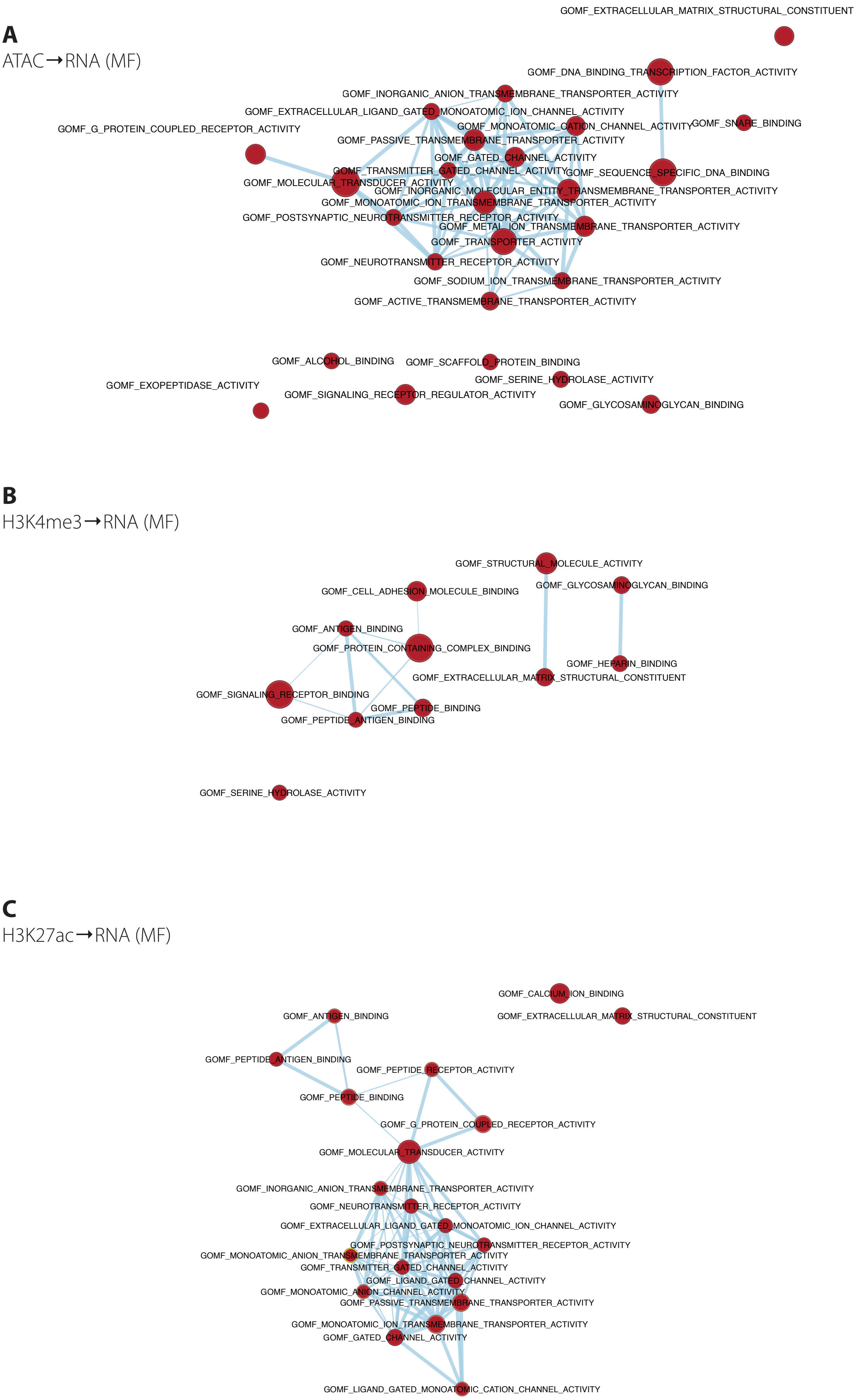


**Supplementary Figure 3. Gene regulatory networks identified by MOBILE.** Pathways identified as significant for GB, with a p-value cutoff of 0.05, for ATAC➞RNA-seq **(A)**, H3K4me3➞RNA-seq **(B)**, and H3K27ac➞RNA-seq **(C)**, for molecular function (MF).

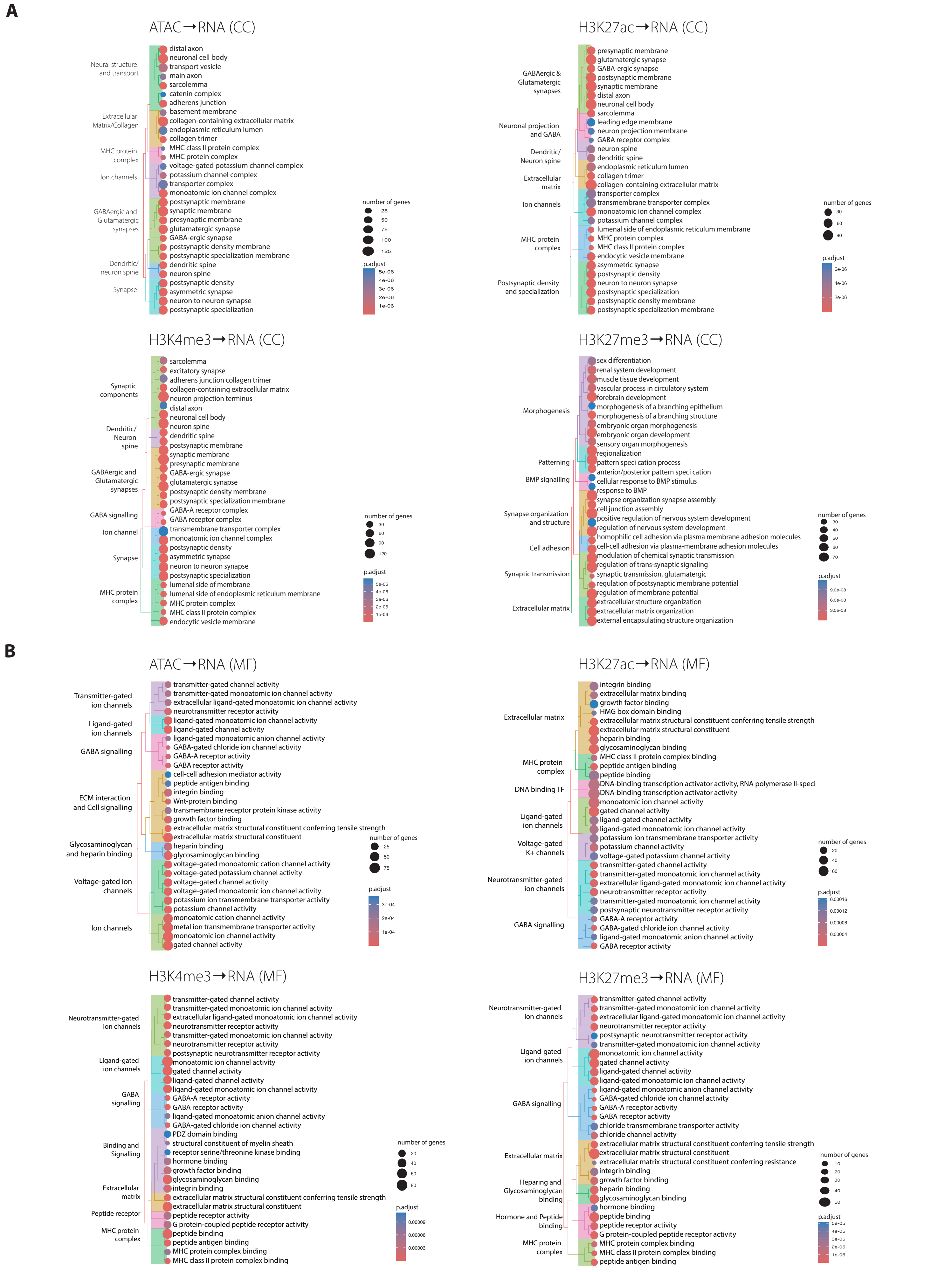


**Supplementary Figure 4. Gene Ontology analysis of the ranked-list genes.** Significantly enriched GO terms for the ranked-list genes from ATAC→RNA , H3K27ac→RNA, H3K4me3→RNA, and H3K27me3→RNA, for **(A)** cellular component (CC) and **(B)** molecular function

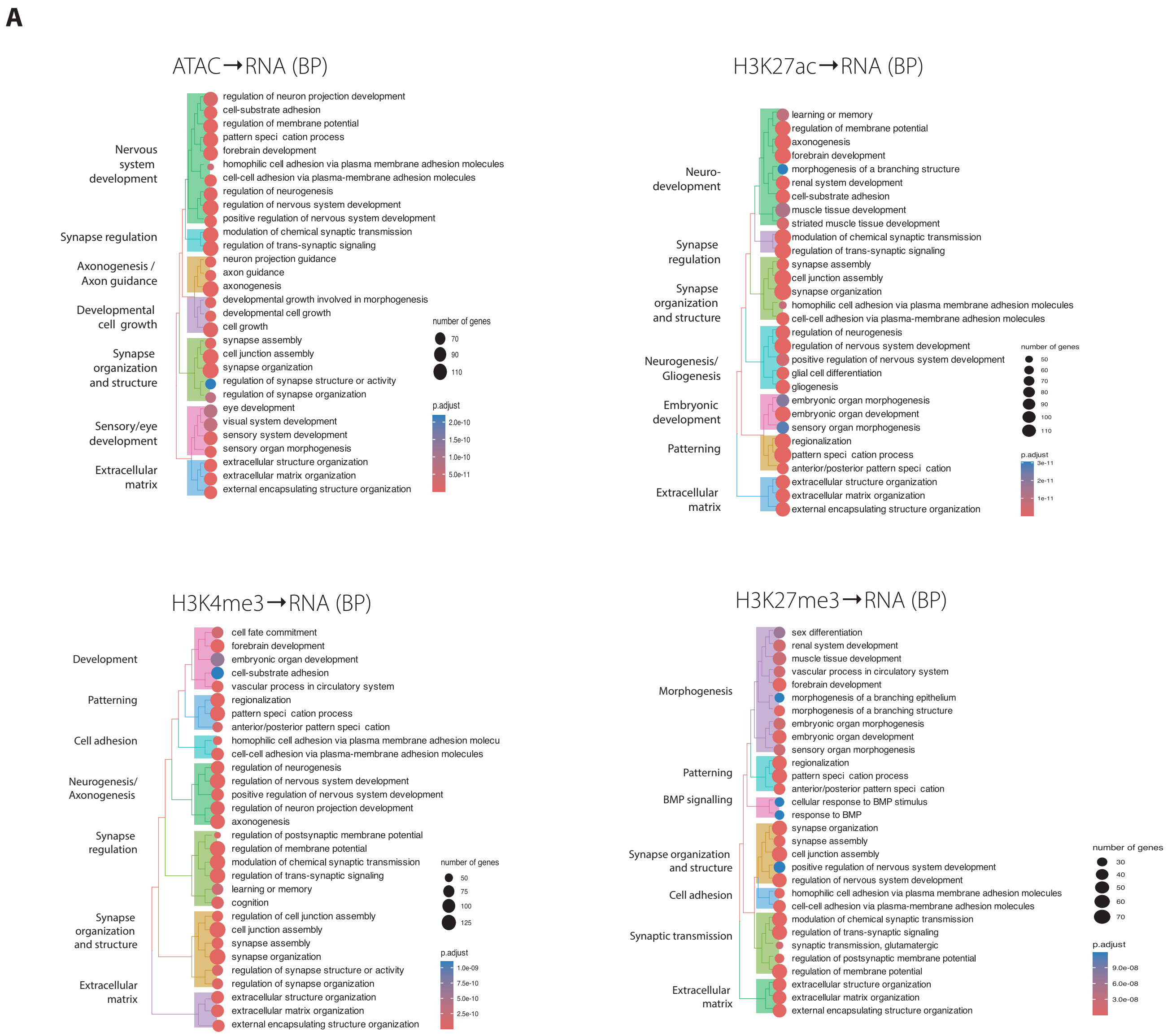


**Supplementary Figure 5. Gene Ontology analysis of the ranked-list genes. (A)** Significantly enriched GO terms for the ranked-list genes from ATAC→RNA , H3K27ac→RNA, H3K4me3→RNA, and H3K27me3→RNA, for biological processes (BP).

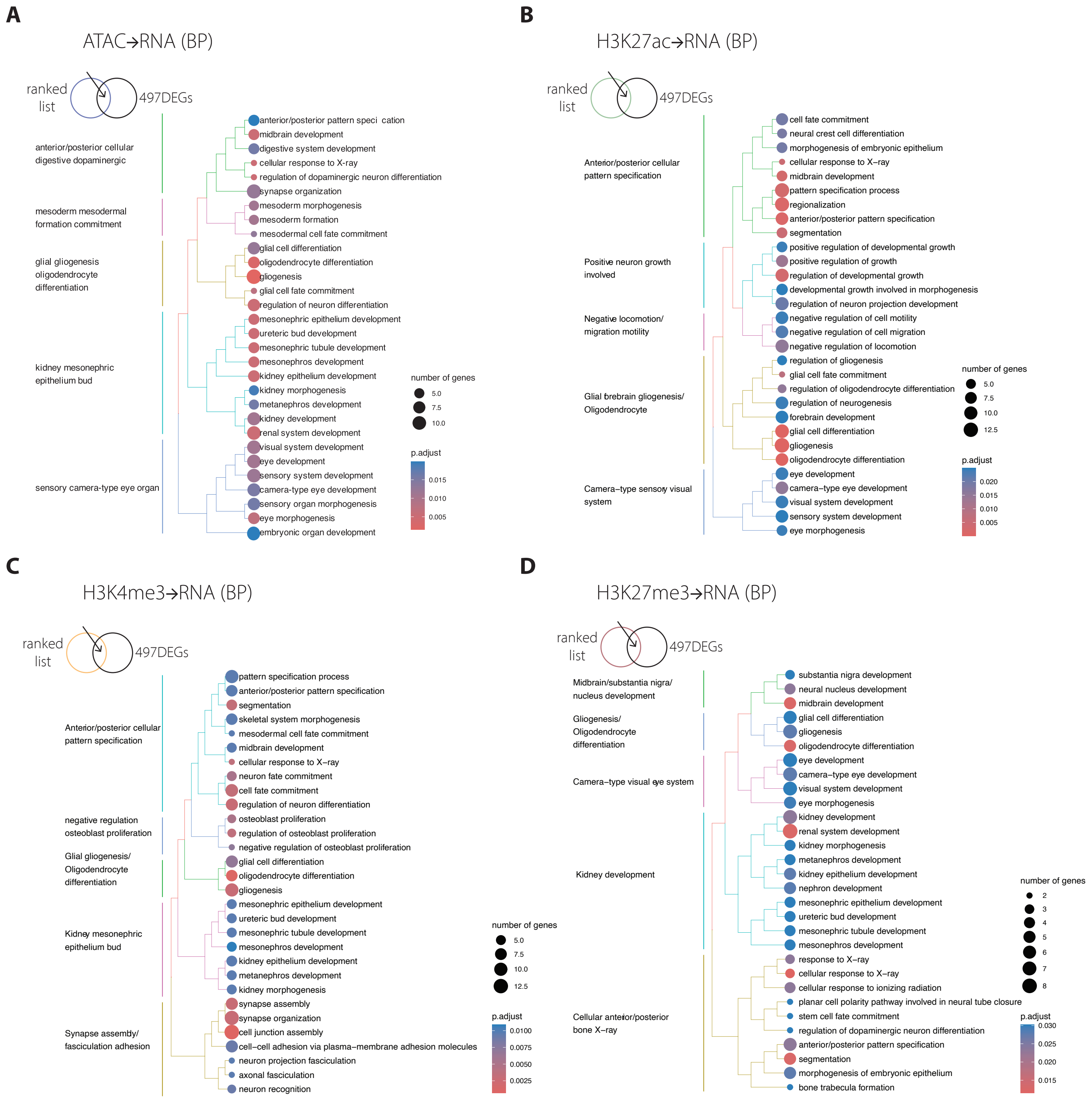


**Supplementary Figure 6. Gene Ontology analysis of the ranked-list genes overlapping with the 497 DEGs from Chakraborty et al., 2023 [7].** Significantly enriched GO terms for 497 DEGs overlapping with the ranked-list genes from ATAC→RNA **(A)**, H3K27ac→RNA **(B)**, H3K4me3→RNA **(C)**, and H3K27me3→RNA **(D)**, for biological processes (BP).

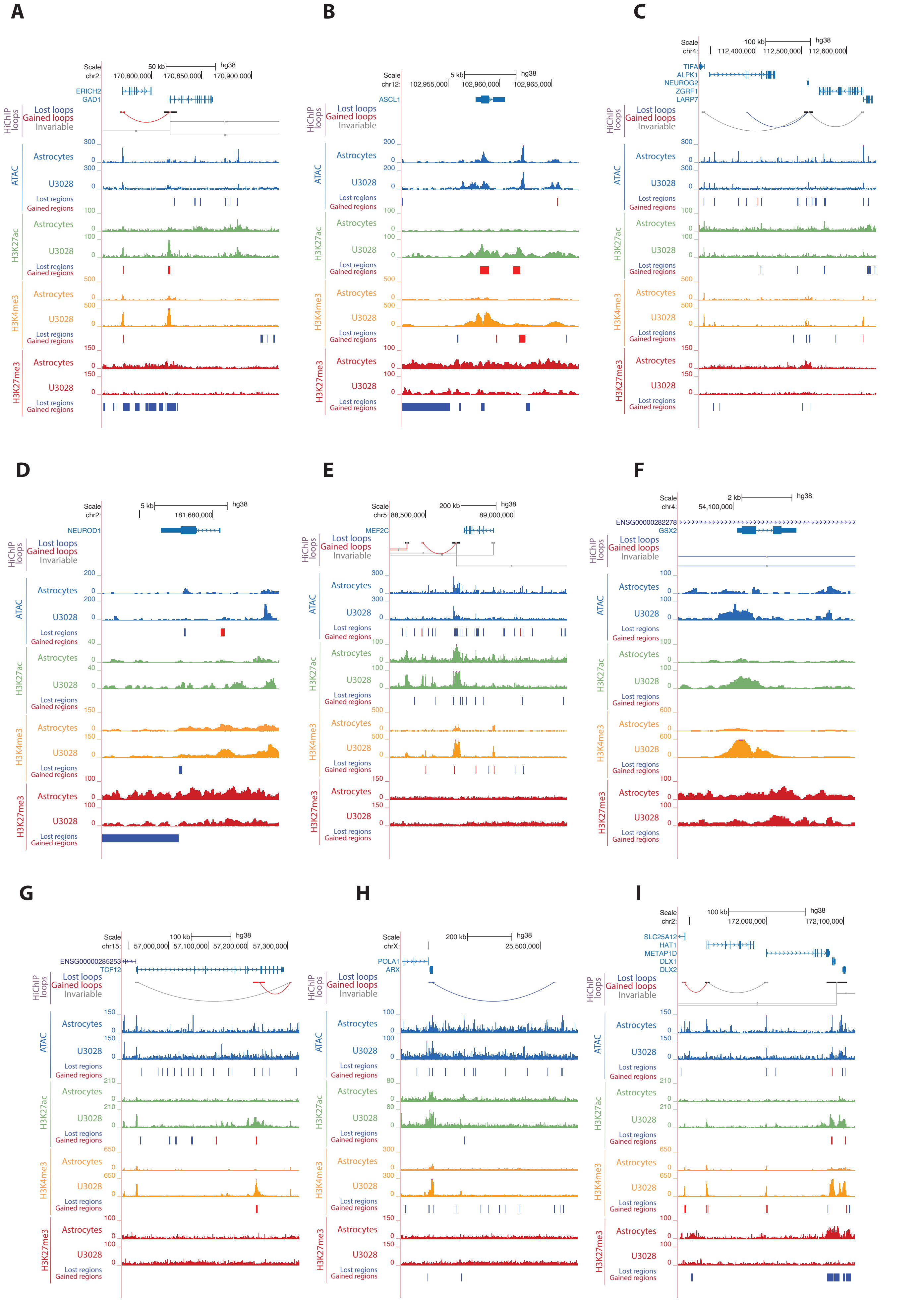


**Supplementary Figure 7. Alterations in the promoter-enhancer interactome and regulatory landscape at the loci of genes related to GABA signaling.** H3K4me3 HiChIP loops, chromatin accessibility (ATAC) and histone modifications (H3K27ac, H3K4me3 and H3K27me3) at the loci of GAD1 **(A)**, ASCL1 **(B)**, NEUROG2 **(C)**, NEUROD1 **(D)**, MEF2C **(E)**, GSX2 **(F)**, TCF12 **(G)**, ARX **(H)**, and DLX1 and DLX2 **(I)**, in normal human astrocytes and one representative patient-derived GB line (Data from Chakraborty et al., 2023 [7]).

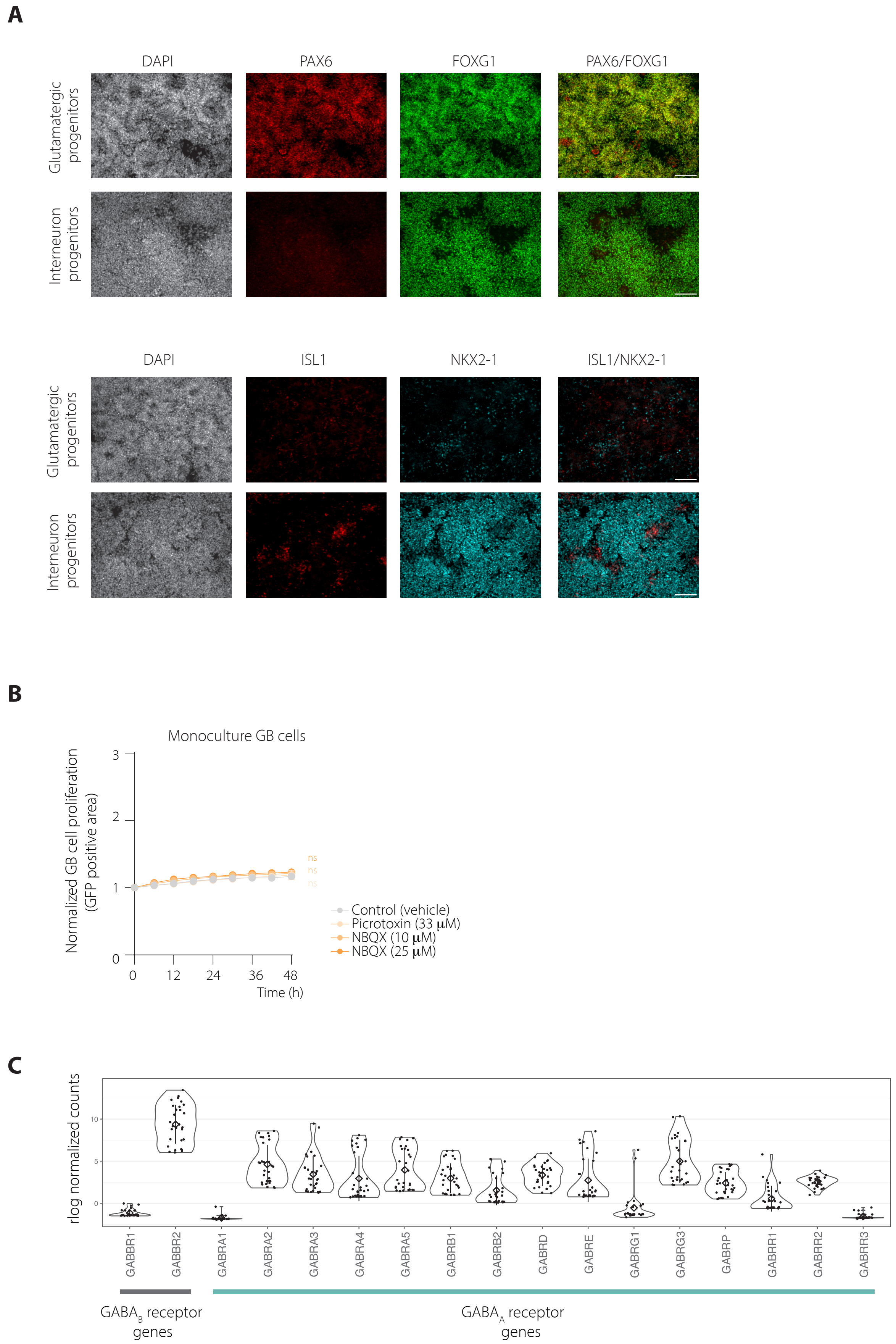


**Supplementary Figure 8. Expression levels of GABA receptor genes in GB cells and specific neural markers in neural progenitor cells. (A)** Immunostaining of glutamatergic progenitors and interneuron progenitors with antibodies against FOXG1, PAX6, ISL1 and NKX2-1 on day 16. Nuclei are counterstained with DAPI. Scalebar=100 µm. **(B)** Proliferation of GB U251-GFP cells over 48 h in monoculture following the indicated treatments [picrotoxin (33µM), NBQX (10µM, 25µM)] compared to the vehicle-control. GB cell proliferation was determined as GFP-positive area, normalized to t=0h and displayed as mean ± SEM. Statistical significance was assessed by unpaired t test with Welch's correction (ns, not significant). **(C)** Expression levels of genes encoding GABA receptors in 15 patient-derived GB lines from Chakraborty et al., 2023 [7].
